# Ecotype Simulation 2: An improved algorithm for efficiently demarcating microbial species from large sequence datasets

**DOI:** 10.1101/2020.02.10.940734

**Authors:** Jason M. Wood, Eric D. Becraft, Daniel Krizanc, Frederick M. Cohan, David M. Ward

## Abstract

**Background:** Microbial systematists have used molecular cutoffs to classify the vast diversity present within a natural microbial community without invoking ecological theory. The use of ecological theory is needed to identify whether or not demarcated groups are the ecologically distinct, fundamental units (ecotypes), necessary for understanding the system. Ecotype Simulation, a Monte-Carlo approach to modeling the evolutionary dynamics of a microbial population based on the Stable Ecotype Model of microbial speciation, has proven useful for finding these fundamental units. For instance, predicted ecotypes of *Synechococcus* forming microbial mats in Yellowstone National Park hot springs, which were previously considered to be a single species based on phenotype, have been shown to be ecologically distinct, with specialization to different temperature and light levels. Unfortunately, development of high-throughput DNA sequencing methods has outpaced the ability of the program to analyze all of the sequence data produced.

**Results:** We developed an improved version of the program called Ecotype Simulation 2, which can rapidly analyze alignments of very large sequence datasets. For instance, while the older version takes days to analyze 200 sequences, the new version can analyze 1.92 × 10^5^ sequences in about six hours. The faster simulation identified similar ecotypes as found with the slower version, but from larger amounts of sequence data.

**Conclusions:** Based on ecological theory, Ecotype Simulation 2 provides a much-needed approach that will help guide microbial ecologists and systematists to the natural, fundamental units of bacterial diversity.

## 1 Background

The identification of closely related, ecologically distinct populations (ecotypes or ecological species) within a microbial community is paramount to understanding the structure and function of the community. Advances in sequencing technology and metagenomic techniques have enabled identification of the major guild-level constituents (e.g., primary producers like *Synechococcus* spp. in hot spring microbial mats) [12, 20]. However, identifying the species-level constituents occupying unique niches within a microbial community can be greatly complicated by the small physical size of the microhabitat and the lack of easily identifiable phenotypic differences amongst closely related but ecologically distinct populations [2].

Early work on microbial species was based on the use of phenotypic differences in cell shape, composition, and metabolism to differentiate between populations. However, systematists were unable to look within natural populations to verify that they were not lumping organisms with similar phenotypes into unnatural groups containing multiple species. With the hope of adding more scientific rigor to species demarcation, molecular divergence cutoffs such as 70% whole-genome DNA-DNA hybridization [43], 95-96% amino acid identity [22], or about 1% identity in 16S rRNA [38] were suggested and calibrated to match the historical phenotype-based groupings. However, little attention was given to whether the resulting sequence clusters came from ecologically distinct populations, and indeed, most named species have been found to include multiple ecologically distinct groups [1, 14, 3, 31, 35, 17, 24, 40, 19, 27, 18, 16]. Cohan and Kopac [4] pointed out that lumping species not only hinders a full understanding of ecological diversity within a bacterial community, but also broadens the apparent diversity within and exaggerates the geographical range of clusters, complicates detection of newly emergent species, and may limit full appreciation of biotechnological potential.

Some have claimed that it would be futile to name and describe all ecological diversity. Doolittle and colleagues have argued that, because of the role of horizontal genetic transfer in bacterial diversification, each individual cell might be ecologically unique and so could be demarcated as its own ecological species [13]. Indeed, in some bacterial groups, particularly the generalist heterotrophs such as *Bacillus*, the rate of bacterial speciation appears to be quite high [23]. Here the discovery of individual ecotypes within a clade may require an extremely highly resolved phylogeny, perhaps based on the entire core genome. On the other hand, speciation appears to be slow in groups with less opportunity for diversification, such as the photoautotrophs, C1-utilizing heterotrophs, and intracellular pathogens [7]. In photoautotrophic, thermophilic, unicellular cyanobacteria of the genus *Synechococcus*, speciation is slow enough that a single gene segment can resolve diversity of ecologically distinct populations [2]. The work we describe here is focused on the slowly speciating *Synechococcus* of Yellowstone hot springs which populate well-established environmental gradients. In such systems, phylogenetically distinct clusters of closely related individuals sharing the same adaptations, metabolic requirements, and susceptibility to the same selection regime exist and can be detected by using a theory-based framework for understanding the diversity.

Fortunately, systematists studying plants and animals have created a long list of species concepts that microbial systematists can utilize for inspiration [41]. Although there are disagreements among these species concepts, most share some common features. That is, each species is a cohesive group (with diversity limited among an ecotype’s members), and each species is founded only once, while different species are ecologically distinct (minimal sharing of resources with other species), and are irreversibly separate [11, 6, 8].

These commonalities in species concepts have been used to formulate multiple models of microbial species. Here we focus on the Stable Ecotype Model in which adaptive evolution within a species is much more frequent than the splitting of species [6]. The Stable Ecotype Model of species and speciation considers the genetic diversity present in a lineage to be the result of two primary variables: net ecotype formation and periodic selection. The rate of net ecotype formation takes into account both extinctions and the formation of new ecotypes. Periodic selection events quash genome-wide diversity within a single ecotype without affecting other ecotypes and can be the result of external forces (e.g., phage infection) or competition within the ecotype with an especially well-adapted mutant or recombinant [6]. In keeping with above-stated criteria, the model defines an ecotype (same as ecological species) to be a cluster of individuals that are ecologically interchangeable, but ecologically distinct from the members of other clusters [21]. Ecotypes so defined have the property of cohesion because the ecological homogeneity within an ecotype allows periodic selection to recurrently purge the ecotype’s diversity [5].

### 1.1 Theory-Based Demarcation of Species Using Ecotype Simulation

We previously developed the Ecotype Simulation algorithm to demarcate ecotypes under the assumption that diversification follows the Stable Ecotype model. It is important to note that the previous algorithm, as well as the present improvement, both use as input only sequence data and no ecological data. Thus, the algorithms allow identification of ecological diversity among closely related populations even before we have characterized the ecological dimensions of diversification. On the way toward ecotype demarcation, the algorithm estimates the essential parameters of the Stable Ecotype model, the net rate of ecotype formation (*omega*) and the rate of periodic selection (*sigma*), as well as the number of ecotypes (*npop*) contained within the sequence sample.

Here we outline the previous Ecotype Simulation algorithm (henceforth termed ES1), previously encoded as a Monte-Carlo style evolutionary simulation program by Koeppel et al. [21]. The ES1 algorithm modeled a lineage of microbes through time as its members speciated and experienced periodic selection events. As shown in Figure 1A, aligned sequence data, which can also be used to generate a neighbor-joining phylogenetic tree using PHYLIP [15], was loaded, processed to remove gaps and PCR errors, then used to construct an *N* × *N* divergence matrix. Complete-linkage clustering was used to obtain the number of sequence clusters (bins) over a range of sequence identity criteria. An example is shown in Figure 2, in which a hypothetical lineage has diversified through time from the last common ancestor (the blue dot) to the extant diversity (labeled A-Q). When the sequence identity criterion was restricted to more closely related sequences, more cluster bins resulted (see Figure 3 and Supplemental Figure 1 for a graphical presentation of “binning curves” generated from environmental sequences). The binning curve was used to estimate the essential parameters of the Stable Ecotype model (the rate of periodic selection, termed *sigma*, and the net ecotype formation rate, termed *omega*, as well as the rate of genetic *drift*, and the number of ecotype populations (termed *npop*). A brute-force method repeatedly simulated the evolution of the lineage using a predefined range of *omega, sigma, drift*, and *npop* values, with the order of events and times at which they occur determined randomly, seeking a combination of values to match the binning curve. A “hill-climbing” stage then utilized the Nelder-Mead simplex method [29] to optimize these estimated *omega, sigma, drift*, and *npop* parameter values through repeated simulations and to calculate confidence intervals for each value.

**Figure 1:**
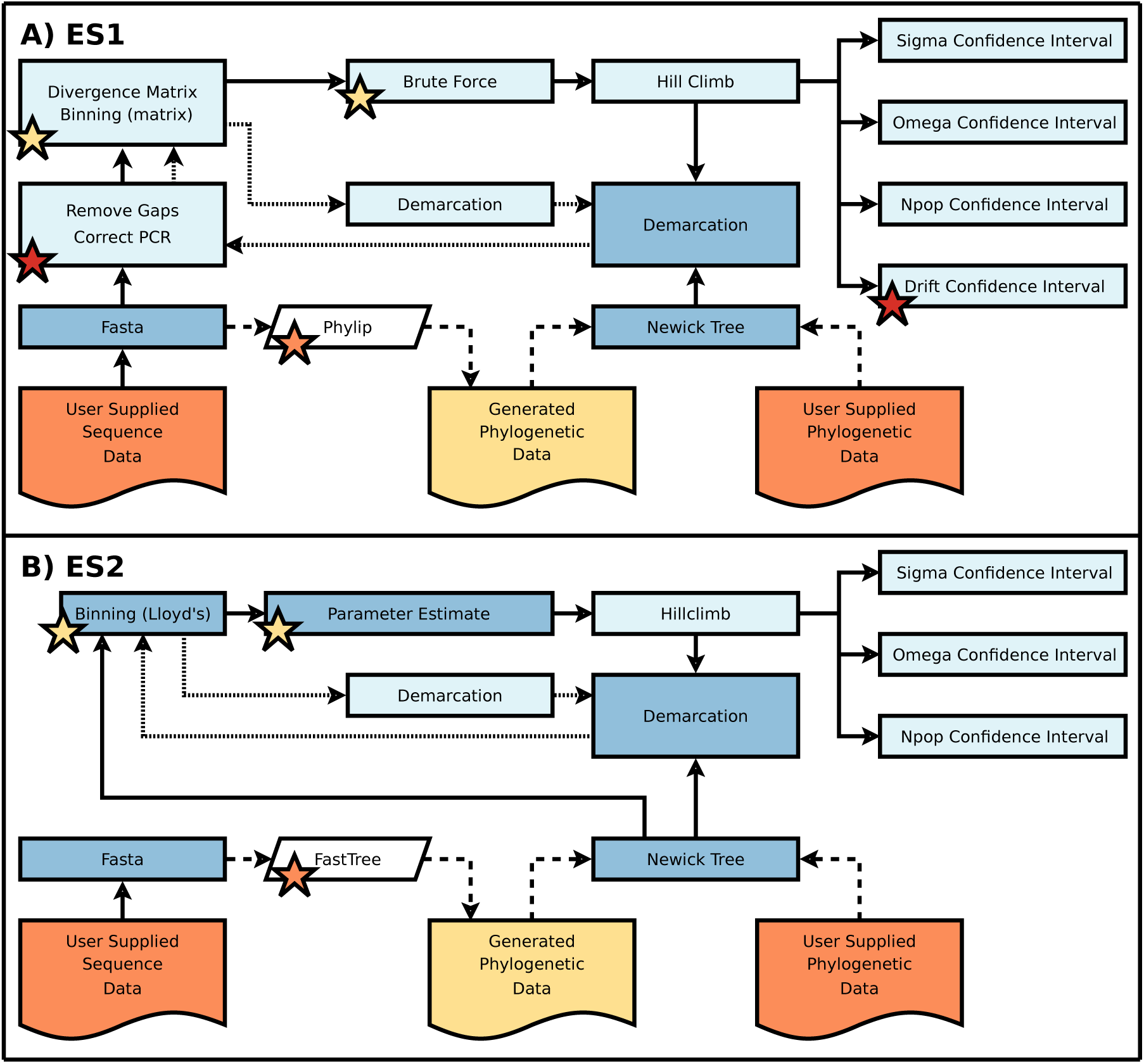
Simplified dependency flowchart of (A) Ecotype Simulation 1 (ES1) and (B) Ecotype Simulation 2 (ES2). Orange nodes are user-supplied data, yellow nodes are generated data, white nodes are external programs, dark blue nodes are Java methods, and light blue nodes are Fortran methods. The dashed lines represent an optional route: the user can either supply phylogenetic data in Newick format, or it can be generated using FastTree from the user-supplied sequence data. The dotted lines represent repeated analyses of various branches of the phylogeny during the ecotype demarcation phase. The red stars in A denote Fortran methods no longer used in ES2. The yellow stars denote Fortran methods in ES1 replaced with Java methods in ES2: Parameter Estimate replaces Brute Force, and the tree-based Binning method replaces Divergence Matrix and matrix-based Binning. The orange stars denote the replacement of PHYLIP in ES1 with FastTree in ES2.

**Figure 2:**
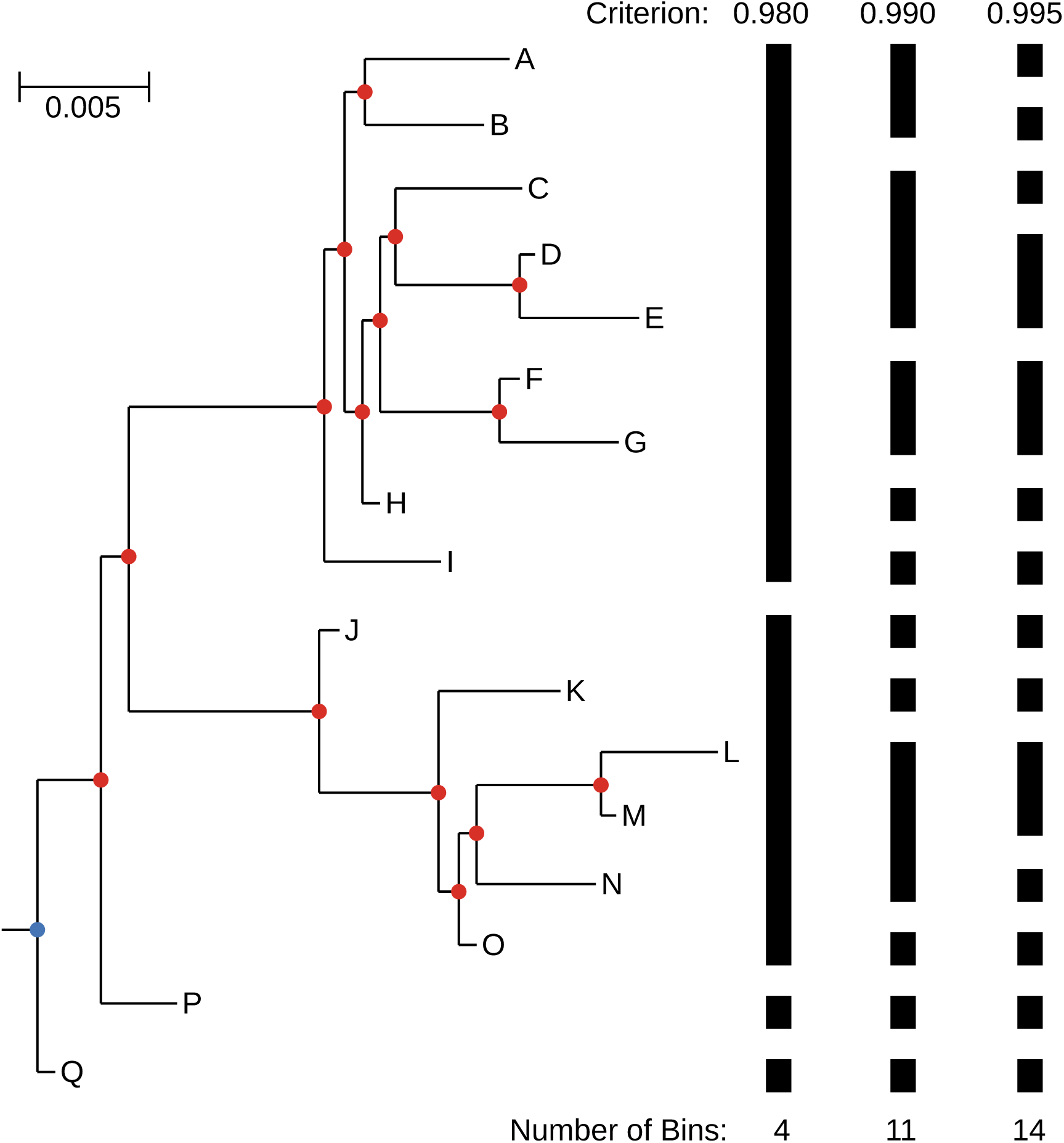
Sequence Identity Criterion Binning of a hypothetical lineage. The root node is marked with a blue dot, internal nodes are marked with red dots, and leaf-nodes that represent individual sequences are marked with the letters A-Q. Black bars denote the result of binning the tree at three different sequence identity criterion values with the number of bins listed below. The scale bar represent 0.005 nucleotide substitutions per site.

**Figure 3:**
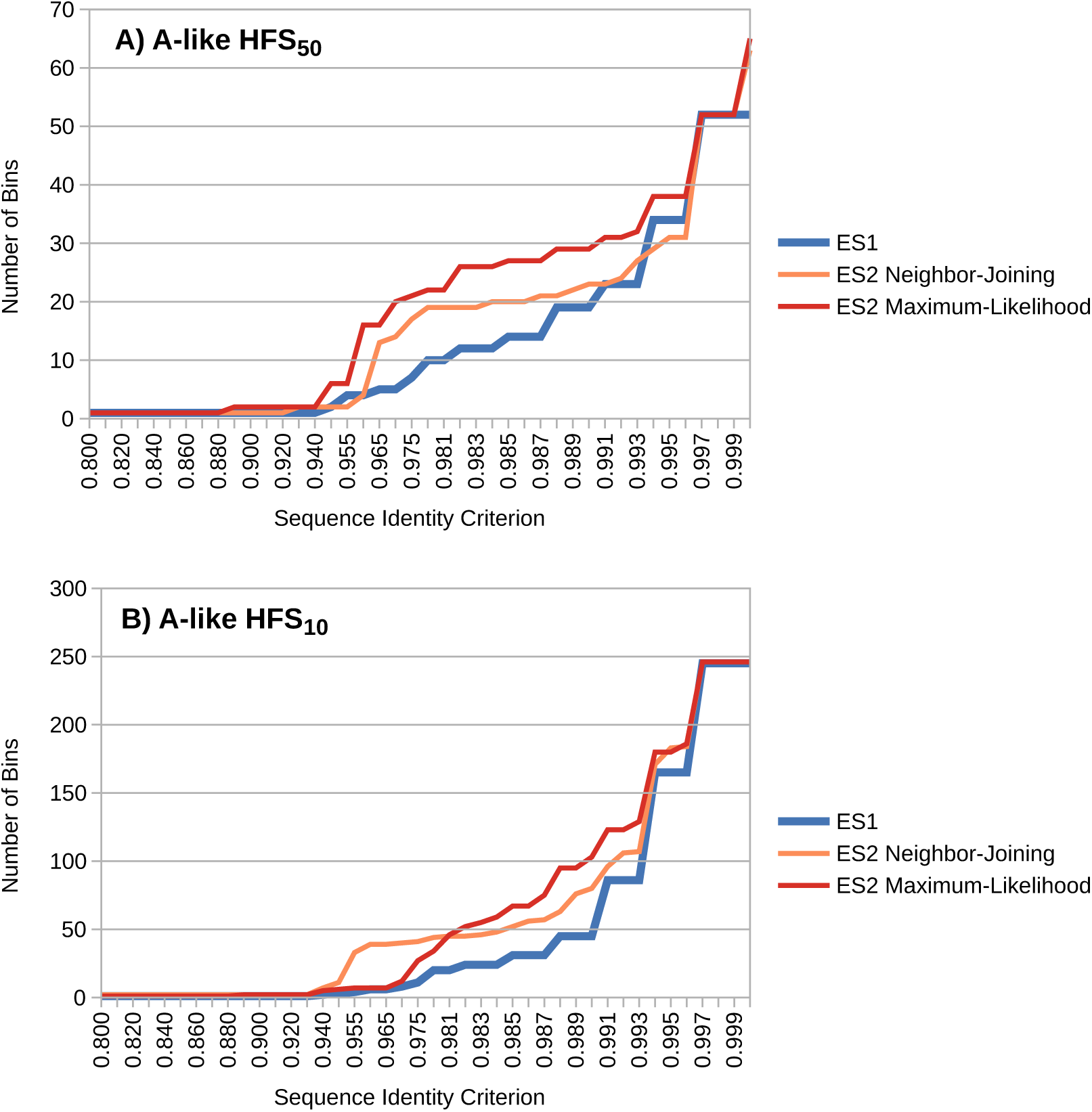
Binning results using A-like *Synechococcus psaA* segment high-frequency sequences (HFSs) occurring at least (A) fifty times (HFS_50_) and (B) ten times (HFS_10_) in the environmental sequence dataset.

The optimized hill-climbing parameter values, along with a phylogenetic tree calculated from the sequence dataset (or provided by the user), are then used in the ecotype demarcation stage, which separately simulates each branch of the tree, starting from the root node (see the blue dot in Figure 2) and progressing though the tree recursively (to the right) until an internal node (red dot) is found with an *npop* value of 1 being within the confidence interval (coarse-scale putative ecotype; PE), or a single leaf node (labeled A-Q) is left, and is demarcated as a PE. Alternatively, fine-scale results [2] require a *npop* value of 1 as the most likely result. The ecotypes predicted by these analyses are tentatively considered PEs until they are proven to have the qualities of ecological species.

### 1.2 Application of Ecotype Simulation to Predict Species in Natural Systems

ES1 has been tested using microbial communities of Yellowstone National Park. A microbial mat community containing the A/B′-lineage *Synechococcus* living along the effluent channel of Mushroom Spring, Yellowstone National Park, WY, USA is known to have closely related 16S rRNA variants differentially distributed along the changing temperature of the flow path ranging from ∼72 °C to ∼50 °C [42]. Importantly, isolates of A-like and B′-like populations, which are found at higher and lower temperatures, respectively, were shown to have different temperature adaptations hypothesized from 16S rRNA distribution patterns. Evidence suggesting ecotypes adaptated to light quantity and/or quality within closely related 16S rRNA lineages required the greater molecular resolution of a faster-evolving gene (*psaA*, which encodes an essential protein subunit of photosystem I) [2]. That is, application of ES1 to the *psaA* segment yielded PEs that were ecologically distinct from one another, as revealed by canonical correspondence analysis (CCA) to differ in their temperature and depth distributions. Moreover, each PE was found to be ecologically homogeneous, in the sense that all members of a group (i.e., individual sequences) were shown to cluster together non-randomly in CCA of temperature and depth distribution. Further evidence of ecological interchangeability emerged from *in situ* experiments that perturbed light and temperature levels. Here, members of the same cluster reacted similarly, but members of different clusters reacted differently to perturbations of light and temperature [2]. These PEs were shown to inhabit different locations within the vertical profile of the mat, with PEs B′9, A1, A4, then A14 progressing from the surface to 1 mm below [2]. More recently, *Synechococcus* isolates that share the same *psaA* sequence as PEs A1 (strain JA-3-3Ab), A4 (strain 65AY6A5), and A14 (strain 60AY4M2) demonstrated different adaptive and acclimative responses to light intensity and quality, with optimal growth *in vitro* similar to conditions present where they can be found in the vertical profile of the mat *in situ* [30]. Comparative genomics and transcriptomics have elucidated genetic differences among these PEs that explain their different metabolic requirements (e.g., low-light adapted PEs were found to have an extra cassette of photosynthetic antenna genes that allow spectral fine tuning under low-light conditions) [34].

Thus, we previously concluded that the hot spring *Synechococcus* PEs demarcated by ES1 hold the essential features of ecotypes – they are ecologically distinct from one another but each is ecologically homogeneous. These analyses provide a well-documented test case for comparing ES1 with the newly developed ES2 algorithm described herein.

ES1 has also been used to study ecotypes of *Bacillus* in the “Evolution Canyons” of Israel and Death Valley. PEs detected by ES1 have been shown to differ in their preferences to solar exposure and soil texture, with membrane differences among PEs potentially explaining heat adaptation differences [21, 9].

### 1.3 The Need for a More Efficient Ecotype Simulation

Although ES1 has proven useful for detecting ecologically distinct clades within groups of bacteria that had previously been considered a single species, the program is extremely slow and is greatly limited by the number of sequences that can be analyzed. Analyzing more than 200 sequences with ES1 can take days on modern computer hardware (Intel Core i7-6700) and then quickly becomes impossible as the number of sequences rises. This is at odds with the large number of DNA sequences that are produced by modern DNA sequencing technologies (Illumina MiSeq can now generate 25 million sequences per run). Various portions of ES1 (Figure 1A) were not designed to manage the large number of sequences produced by modern sequencing technologies (e.g., the memory usage of the ES1 matrix-based binning algorithm increases with the square of the number of sequences and the CPU usage increases with the cube of the number of sequences). The phylogenetic algorithms provided by PHYLIP and used by ES1 suffer from the same problem, and would offer no help in analyzing the large number of sequences produced with modern sequencing techniques. Other parts of the ES1 program needlessly store excess data in memory (e.g., both the Java object that loads Fasta-formatted sequence data and the Fortran simulation code store all sequences in memory even though the sequence data are not used during the simulation). By addressing these and other issues in this new version of Ecotype Simulation (henceforth termed ES2), we show that it is possible to successfully predict ecological species from very large datasets within a few hours.

## 2 Implementation

### 2.1 Overview of Changes to Ecotype Simulation

ES1 and ES2 work in roughly the same way (see Figures 1A and B for an overview comparison) with some fundamental differences that may affect the results. ES2 has a redesigned graphical user interface that will be familiar to ES1 users but is more functional (Figure 4). ES2 also has a new command line interface for use when a graphics interface is not available. Where ES1 uses a phylogenetic tree only during the demarcation stage, ES2 uses a phylogenetic tree for all stages of the analysis. ES2 incorporates FastTree [36], rather than treeing algorithms found in PHYLIP [15], if the user wishes to use ES2 to generate the phylogenetic tree.

**Figure 4:**
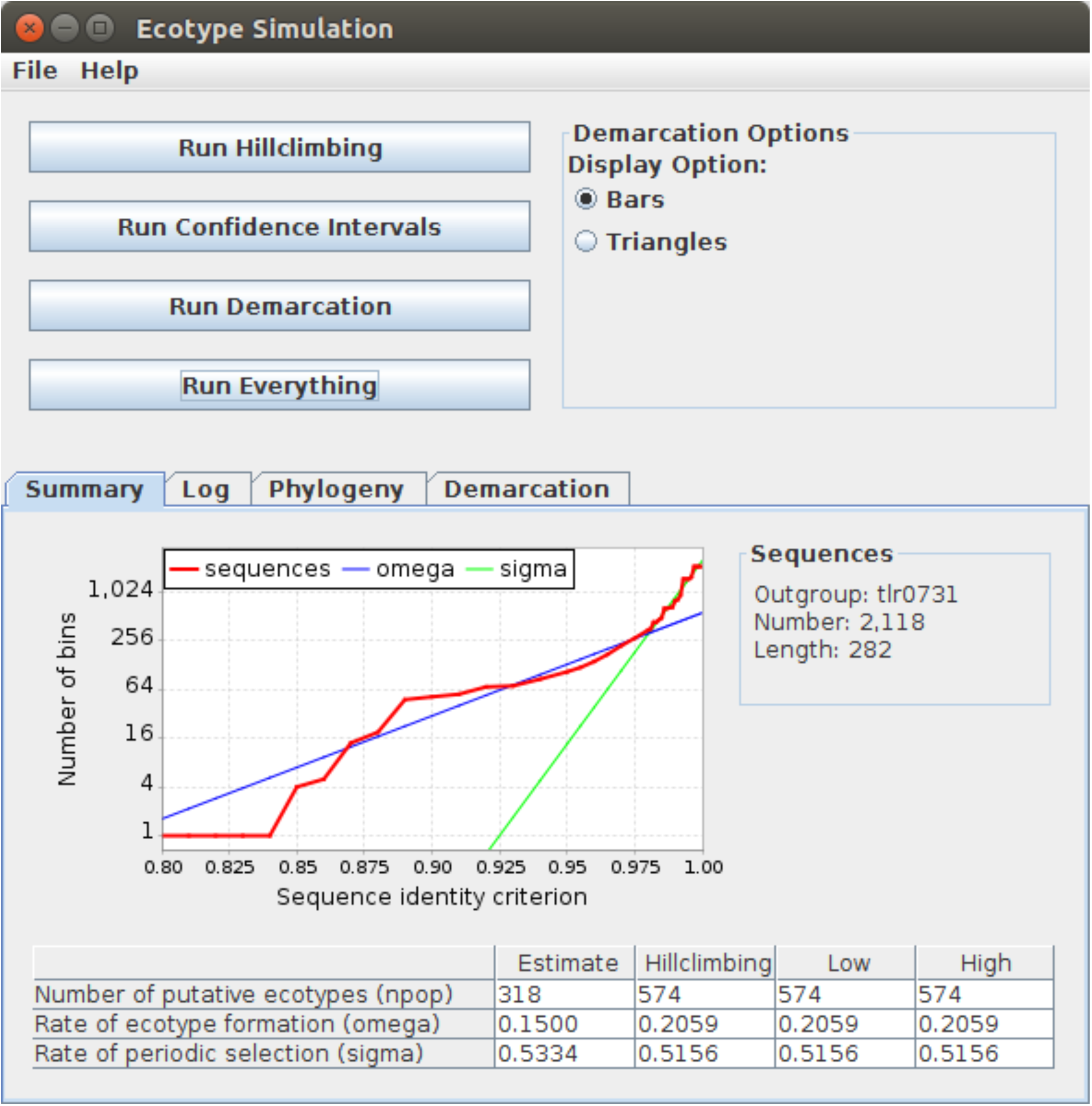
Screenshot of the Ecotype Simulation 2 graphical user interface, with the Summary tab displayed.

ES1 and ES2 both calculate the number of sequence clusters (bins) to quantitatively summarize the diversity present in a sequence dataset at various levels of sequence identity criteria, but ES2 utilizes a new tree-based binning method that avoids the need to clean up the sequence data (beyond what is necessary to produce a tree) or to spend considerable time calculating a divergence matrix (see Section 2.1.2 and Figure 2 for an overview and Figure 3 for a comparison in resulting curves). Both ES1 and ES2 then use these bins to find initial estimates for the rates of periodic selection (*sigma*) and net ecotype formation (*omega*), and to estimate the number of ecotype populations (*npop*). However, ES2 uses a custom version of Lloyd’s algorithm [26] for the initial parameter estimation that converges on a likely solution much faster than the ES1 brute-force method. ES1 also estimates the rate of genetic drift, but since population sizes of bacteria are huge [37], genetic drift can be safely ignored and has been removed from the simulation in ES2. In both ES1 and ES2, these initial estimates are then used as the starting point in a hill-climbing stage to find the maximum-likelihood estimates for the three parameters. In the ES2 hill-climbing stage, the estimation of *omega, sigma*, and *npop* confidence intervals, and the ecotype demarcation stage are all simulated in the same manner described above for ES1, but with some fundamental differences and numerous optimizations made to the simulation that will be described below.

#### 2.1.1 Phylogenetic Trees

To generate phylogenetic trees from sequence data for the binning and ecotype demarcation stages of ES2, FastTree [36] has been incorporated into the program to generate approximate maximum-likelihood trees. FastTree is able to perform single-gene or whole-genome sequence alignments using less memory than distance-matrix based methods. That is, memory requirements for FastTree are proportional to [*N* × *Length* × 4 + *N* ^1.5^], while distance-matrix based methods are proportional to [*N* × *Length* × 4]^2^, with N being the number of sequences. FastTree also requires much less CPU time (proportional to [*N*^1.5^ × *log*(*N*) × *Length* × 4]) than distance-matrix based methods (proportional to [*N* × *Length* × 4]^3^), and provides higher topological accuracy than distance-matrix based methods [36]. Since the ES1 binning method utilizes a distance-matrix and performs operations similar to the neighbor-joining method, we expect the higher topological accuracy of FastTree to benefit the new ES2 tree-based binning method.

#### 2.1.2 Binning

ES2 derives its complete-linkage binning solution for each level of sequence identity directly from the tree, without the need for calculating an *N* × *N* matrix of pairwise divergences. Whereas the ES1 matrix-based algorithm computes the distance between all members of the sequence dataset (i.e., all possible pairs of sequences, *N* ^2^), the new tree-based algorithm of ES2 computes only the maximum distance among leaf-nodes that represent each sequence in the dataset (see the labels A-Q in Figure 2) and the internal nodes that connect them together (red and blue dots). For instance, in Figure 2, sequences A and B both exceed an identity criterion of 0.995 and are demarcated as separate bins. At an identity criterion of 0.990, they are combined into a single bin, and at an identity criterion of 0.980, they are grouped with sequences C-I into a much larger bin. But, once defined, each sequence or bin is ‘pruned’ from the tree. Thus, by using a tree-based algorithm, ES2 is able to avoid calculating distances between nodes that share no direct relation, which offers two primary advantages. First, ES2 is able to calculate the number of bins using a method that scales linearly with the number of sequences rather than the ES1 matrix-based binning method that scales cubically. Second, ES2 can now use binning results during all stages of the program that were calculated using the same phylogeny that ES1 used only in the final ecotype demarcation stage.

#### 2.1.3 Estimating *omega, sigma*, and *npop*

The algorithm that estimates the initial values of *omega, sigma*, and *npop* for the hill-climbing stage in ES1 (Figure 1) used a brute-force approach – it searched a large range of predefined *omega, sigma*, and *npop* parameter values for likely results. A faster approach was needed to achieve the goals established for ES2. The Stable Ecotype model predicts that sequence variation is divided into two phases: a phase responsible for the formation of the ecotypes and a phase of random mutations that have occurred after the last periodic selection event. This is reflected in the binning data. During the initial phase, bins are created at a steady rate representing the creation of new ecoptyes. Once the second phase is entered the number of bins rises sharply until each unique sequence forms its own bin. Using a custom variant of Lloyd’s k-means algorithm [26] designed to partition point data into clusters based on distance from a line rather than from a point, the ES2 approach fits two lines to the base-two logarithm of the binning data in order to capture these two phases. The graph in Figure 4 shows the two fitted lines. The slope of the blue line estimates the rate of ecotype formation – *omega*. The number of bins at the intersection of these two lines serves as the estimate for *npop* – the number of predicted ecotypes. The binning criteria at which this intersection occurs provides an estimate of the time since the last periodic selection, thus providing an estimate of *sigma* – the rate of periodic selection. Once these values have been estimated, ES2 proceeds as in ES1 to apply hill-climbing with these values as a starting point to find a maximum-likelihood estimate for the parameters.

#### 2.1.4 The Simulation

Multiple changes were made to the simulation code used by hill-climbing, demarcation, and the *omega, sigma*, and *npop* confidence intervals to decrease its runtime and memory usage. First, and most vital, the simulation now uses an ultrametric approach to estimate the number of mutations in each internode, rather than drawing from a Poisson distribution. Second, as mentioned above, the rate of genetic drift is no longer calculated or used in the simulation. Third, the waiting time to an event (either ecotype formation or periodic selection) was previously estimated through simulation-style code in ES1, but in ES2, the waiting time to a “key event” is estimated using a mathematical function (*timeWait* = −1 × *log*(*x*)*/rateKey*, with *x* chosen randomly from a uniform distribution in the range [0, 1)) that runs in constant time while retaining the stochasticity of *timeWait*. The rate of key events (*rateKey*) is the sum of the rates *omega* and *sigma*. All three changes make the overall simulation much faster without greatly affecting the results (see Section 3.2).

To allow ES2 to fully utilize modern multi-core computers, OpenMP [10] was used to thread the main-loop of the simulation that is repeatedly run during the course of the simulation. To overcome an inability to use the intrinsic Fortran 90 pseudo-random number generator in a threaded environment, the Ziggurat algorithm [28], was translated from C source code into Fortran 90, and added to ES2. The Ziggurat algorithm was modified to use an object-like interface to store the state variables for the pseudo-random number generator, enabling a separate state to be used for each thread.

One last noteworthy change to the simulation involves the addition of a new Fortran 90 module to store dynamically sized arrays. ES1 used statically sized arrays to store variably sized content, and was usually successful at avoiding stack-overflow errors by using overly large default values for the array size. However, no error checking was done in ES1, and the program would occasionally fail with cryptic error messages. This new module dynamically increases the size of the array as needed, allowing ES2 to store variably sized content and fixing ES1 stability issues.

#### 2.1.5 Ecotype Demarcation

The ecotype demarcation stage has been slightly simplified in order to increase its speed. For a given subclade, the ES1 version of the algorithm tested all *npop* values between 1 and the current *npop* estimate for the entire clade to find the most likely number of ecotypes within the subclade. Since the point of this stage is to determine whether the subclade currently being examined makes up a single or multiple ecotypes, ES2 has been simplified to assume that *npop* > 1 if the average success rate of 1000 trials is found to be zero when *npop* = 1 for the subclade. Only when the average success rate of these trials with *npop* = 1 is greater than zero are other values of *npop* tested using a likelihood-ratio test for whether they are significantly more likely (*α* = 0.05). The best case scenario for this optimization results in the ecotype demarcation stage only testing a single *npop* value for a subclade in the phylogeny. The worst case scenario matches the ES1 behavior of testing every *npop* value between 1 and *npop* for a subclade.

### 2.2 Evaluation of Ecotype Simulation

#### 2.2.1 Environmental Sequences

Because of the slow speed of ES1, Becraft et al. [2] had to restrict analysis to high-abundance environmental sequence segments, defined as those with >50 identical copies across the entire dataset (HFS_50_). There were 65 A-like *Synechococcus psaA* genotypes, and 88 B′-like genotypes in the HFS_50_ database. To compare the ability of ES2 to predict ecotypes inhabiting the Mushroom Spring microbial mat predicted by ES1, we used the same set of sequences. To demonstrate the enhanced capabilities of ES2, we used sequences with >10 identical copies across the entire dataset (HFS_10_). In the HFS_10_ analysis, there were 246 A-like *Synechococcus psaA* genotypes and 298 B′-like genotypes. The script that Becraft et al. [2] used to find the HFS_50_ was used to find the HFS_10_ from the same environmental sequence dataset (`hfs-finder.pl`; available from https://github.com/sandain/pigeon). Each unique HFS was a single entry in each analysis. In the main text, we focus on predominant PEs in the B′-lineage (PE B′9) and A-lineage (PEs A1, A4, and A14) *Synechococcus*, which were found in ES1 analysis to have distinct vertical positioning in the mat at 60 °C to 63 °C. We use PE B′9 and the A-lineage PEs as examples in the main text, but include similar analyses for the remaining B′-lineage in the Supplemental Information (Supplemental Figures 2, 3, 4, and 5).

#### 2.2.2 Tree Building

Because ES2 substituted the maximum-likelihood phylogenetic trees generated by FastTree for neighbor-joining phylogenetic trees generated by PHYLIP used in ES1, we needed to compare analyses made with both tree building algorithms. The neighbor-joining trees used by Becraft et al. [2] for analyses of the *Synechococcus* A- and B′-like HFS_50_ subset of sequences were used here for comparison (Figure 5 and Supplemental Figure 2). To generate similar neighbor-joining trees for the HFS_10_ subset of sequences, the combination of PHYLIP’s `dnadist` and `neighbor` programs [15] were used with default parameters. FastTree [36] was used to build approximate maximum-likelihood trees for both the HFS_50_ and HFS_10_ subsets of sequences (Figure 6 and Supplemental Figure 3). PEs were separately demarcated for A- or B′-like *Synechococcus* neighbor-joining or maximum-likelihood phylogenies, generated from HFS_50_ or HFS_10_ datasets.

**Figure 5:**
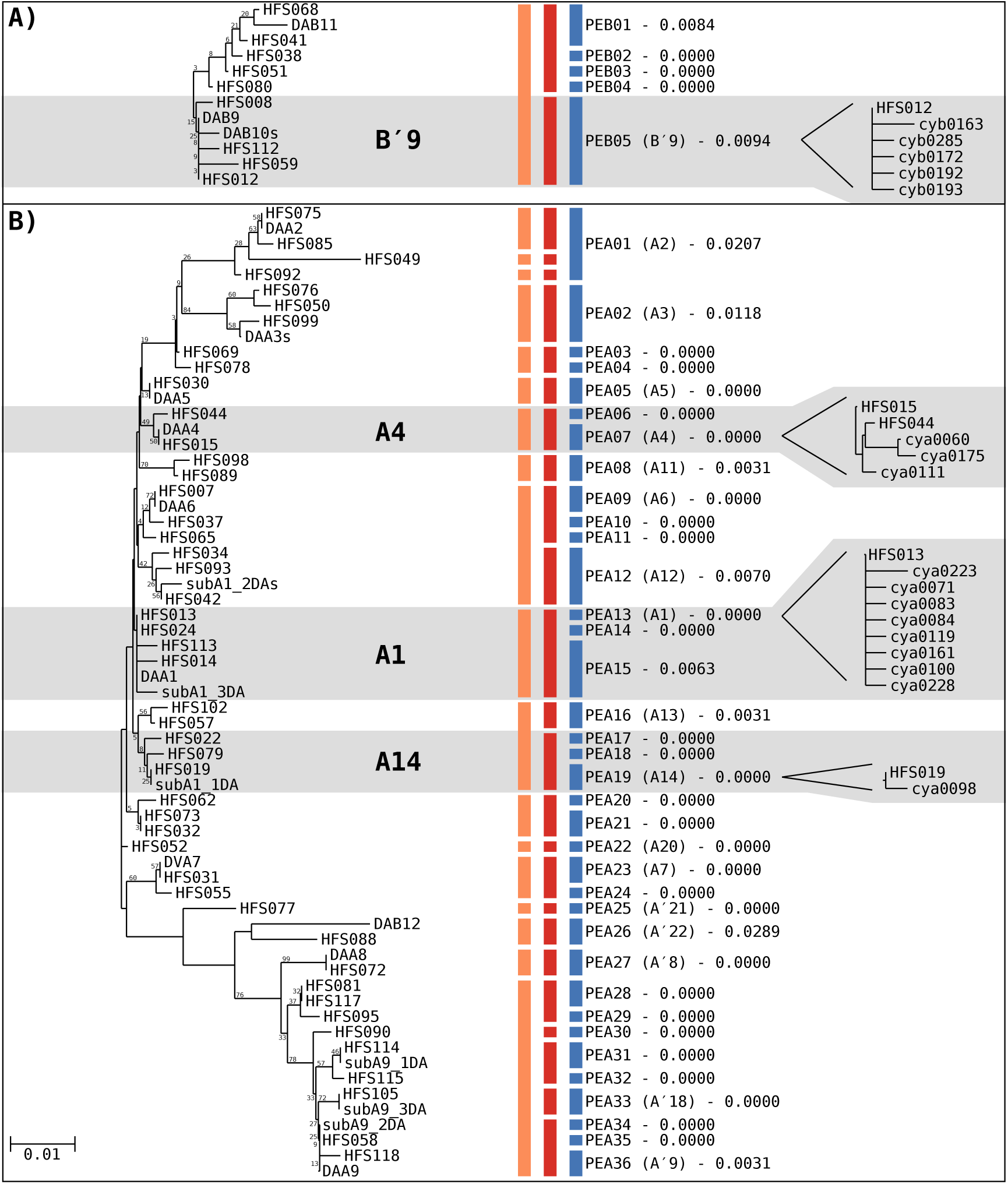
Neighbor-joining phylogeny with putative ecotype (PE) demarcation of (A) PE B′9- and (B) A-like *Synechococcus* HFS_50_ environmental *psaA* segments generated using PHYLIP. Gray shading denotes predominant PEs demarcated using Ecotype Simulation 1 (ES1) that are examined in detail in the main text. Colored vertical bars indicate demarcation done by different algorithms, from left to right: orange, ES1 coarse-scale; red, ES1 fine-scale; and blue, Ecotype Simulation 2 (ES2). PEs are labeled with the ES2 demarcation and include the maximum distance among members of the clade. ES2-demarcated PEs that contain the same dominant variant as a PE demarcated by Becraft et al. [2] using ES1 are indicated in parentheses after the ES2-generated PE names. Portions of the HFS_10_ neighbor-joining tree corresponding to predominant PEs PEB05 (B′9), PEA07 (A4), PEA13 (A1), and PEA19 (A14) are included on the right for comparison. The scale bar represent 0.01 nucleotide substitutions per site and is the same in parts A and B.

**Figure 6:**
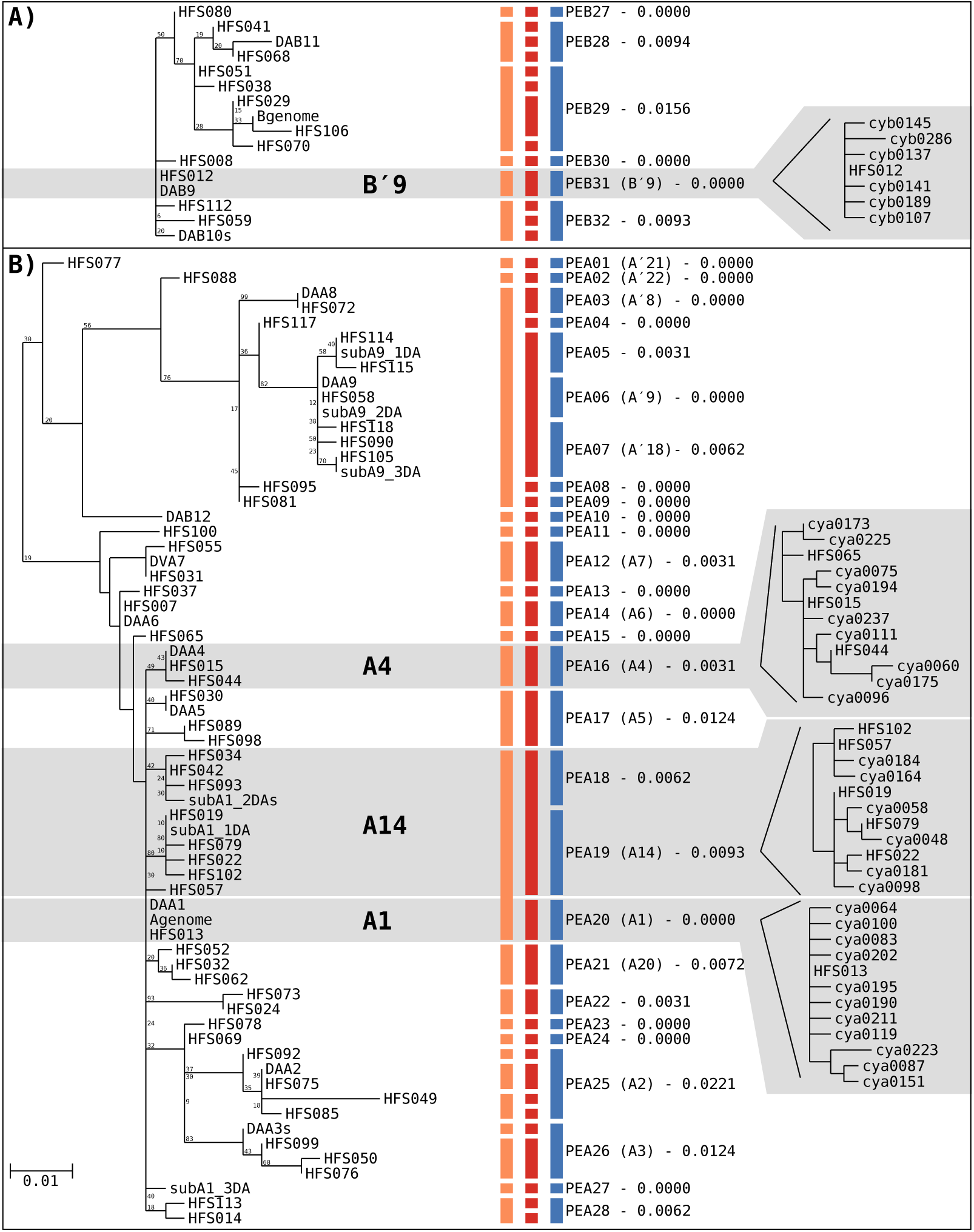
Maximum-likelihood phylogeny with putative ecotype (PE) demarcation of (A) PE B′9- and (B) A-like *Synechococcus* HFS_50_ environmental *psaA* segments generated using FastTree. Gray shading denotes predominant PEs demarcated using Ecotype Simulation 1 (ES1) that are examined in detail in the main text. Colored vertical bars indicate demarcation done by different algorithms, from left to right: orange, ES1 coarse-scale; red, ES1 fine-scale; and blue, Ecotype Simulation 2 (ES2). PEs are labeled with the ES2 demarcation and include the maximum distance among members of the clade. ES2-demarcated PEs that contain the same dominant variant as a PE demarcated by Becraft et al. [2] using ES1 are indicated in parentheses after the ES2-generated PE names. Portions of the HFS_10_ maximum-likelihood tree corresponding to predominant PEs PEB31 (B′9), PEA16 (A4), PEA19 (A14), and PEA20 (A1) are included on the right for comparison. The scale bar represent 0.01 nucleotide substitutions per site and is the same in parts A and B.

PEs demarcated by ES2 from A- or B′-like *Synechococcus* sequences were named using the pattern PEAxx or PEBxx, respectively (with xx being a number between 01 and 36, the maximum number of PEs detected in our analyses). Since clade structure changes between tree algorithms and datasets used, PE names assigned by ES2 in different analyses do not usually correspond. Furthermore, PEs newly demarcated and named by ES2 analyses do not have names that correspond with previously named PEs demarcated with ES1 by Becraft et al. [2]. We identified the corresponding ES2-demarcated PE by the presence of the most dominant variant in the ES1-demarcated PE of Becraft et al. [2]. This correspondence is indicated by identifying the PE previously demarcated by ES1 in parenthesis following the corresponding ES2-demarcated PE assignment. For instance, the ES1 PE A1 demarcation by Becraft et al. [2] (both fine- and coarse-scale) was split by ES2 into PEs PEA13 (A1), PEA14, and PEA15 in neighbor-joining analysis (Figure 5), but only PEA13 (A1) retains the original PE designation, since it contains the same dominant variant (HFS013) as the ES1 PE A1.

#### 2.2.3 Canonical Correspondence Analysis

Canonical correspondence analysis (CCA) [39, 25] provided by the `R` package `vegan` [32] and a new version of the custom plotting software (`cca.R`; available from https://github.com/sandain/R) that was used by Becraft et al. [2] were used to compare the ecological distribution of the members of PEs with the sampled environmental parameters (temperature of the water and depth within the mat). The community data matrix analyzed by CCA was created with the `hfs-counter.pl` script (available from https://github.com/sandain/pigeon) that simply counts the abundance of each HFS in each environmental sample. Although PEs were demarcated separately for A- and B′-like *Synechococcus* variants, the CCA community data matrix was comprised of variants from both lineages.

#### 2.2.4 Testing for Ecological Interchangeability and Ecological Distinctness

Canonical correspondence analyses provide a method for testing the ecological inter-changeability and ecological distinctness of a single PE. Ecologically interchangeable members of a PE are expected to form non-randomly distributed clusters of sequences within the ordination space of the CCA. The plotting software mentioned above reports a p-value for each PE demarcation that represents the probability that the observed distribution of the PE in ordination space is different from random. Ecologically distinct PEs are expected to form clusters separate from other PEs in the CCA ordination space. Deviation from either of these expectations does not mean that the ecotype demarcation is incorrect, because PEs that overlap in ordination space could be a result of the niche-defining variable that differentiates the populations having not been measured. A cluster with a distribution that cannot be differentiated from random could be the result of limited sampling of the PE (i.e., a PE with only two members may appear to be randomly distributed while a PE with the exact same spread in the ordination space but with fifty members may appear distributed differently from random). The researcher must examine each PE predicted by ES2 for these conditions (e.g., limiting the analysis to sequence types with abundances >10 to limit the influence of undersampling) and determine whether they have been satisfied before promoting the putative ecotype to an ecotype. Care must still be taken when interpreting PE distributions with CCA because ES2 may demarcate PEs using a gene segment shared between closely related populations that have otherwise diverged in the environment. Ecological species, or ecotypes, should be thought of as the smallest clades meeting the expectations of ecological distinction and ecological interchangeability. Although PEs with only a single member are included in CCA analyses, it is impossible to test for ecological interchangeability in such PEs.

#### 2.2.5 Analysis of ES2 Runtime

To demonstrate the power of ES2, we analyzed a dataset containing 2,197,037 A-like *Synechococcus psaA* segments obtained from the Mushroom Spring mat using Illumina MiSeq [44]. To test the capabilities and results of ES2 using variously sized sequence datasets, the entire set of 191,901 unique sequences representing this dataset was randomly subsampled every 5,000 sequences from 5,000 to 190,000 sequences (i.e., 5,000, 10,000, .., 190,000; see below) with the `unique random fasta.pl` script (available from https://github.com/sandain/pigeon) and a maximum-likelihood tree was generated with FastTree. Each subset of sequences and its pre-generated tree was run with the ES2 command line interface three times for replication on an Intel Core i7-6700 processor running Ubuntu 17.10, and timed using GNU `time` version 1.7.

We also analyzed a dataset containing 1,774,277 B′-like *Synechococcus psaA* segments representing 195,153 unique sequence types in a similar manner (see Supplemental Figure 5).

## 3 Results and Discussion

### 3.1 Binning

Here we examine the efficacy of the new tree-based binning method of ES2 compared to the distance-matrix-based binning method of ES1. We generated neighbor-joining trees using PHYLIP and maximum-likelihood trees using FastTree from the same sequence data binned previously using ES1. The neighbor-joining method provided by PHYLIP utilizes a matrix-based method that is very similar to the ES1 matrix-based binning method. Thus, it is not surprising that the ES2 binning results from neighbor-joining trees more closely follow the ES1 binning results of the environmental sequences than the ES2 results from the maximum-likelihood tree (compare the orange and red lines to the blue lines in Figure 3 and Supplemental Figure 1).

### 3.2 Performance of ES1 and ES2

Figures 5 and 6 are neighbor-joining and maximum-likelihood phylogenies for variants comprising PE B′9 and all A-like PEs. PEs are numbered according to ES2 output, and correspondence with predominant PEs predicted using ES1 by Becraft et al. [2] is shown in parentheses. Although these trees look different, for example the A′ PE cluster found at that bottom of the neighbor joining tree (Figure 5B) is located at the top of the maximum likelihood tree (Figure 6B), careful inspection reveals that there are only rare cases in which individual HFS variants did not cluster similarly in these two trees.

As previously noted in that study, fine-scale ES1 analysis yielded a greater number of PEs than coarse-scale analysis, but this was mainly due to the effect of the different demarcation rules used in fine-scale analyses splitting coarse-scale demarcated PEs. For instance, in the fine-scale analyses shown in Figures 5 and 6, 51 to 57% of the PEs were identical to those in coarse-scale analysis and 42 to 49% resulted from splitting. ES2 analysis yielded a greater number of PEs, and these differences were mainly due to splitting ES1 coarse-scale PEs into multiple PEs. In the neighbor-joining tree (Figure 5) 20% of the PEs demarcated by ES2 were identical to those demarcated by coarse-scale ES1 analysis, with 78% of the new PEs resulting from splitting, and only 2% of the PEs were due to lumping. Similarly, in the maximum likelihood analysis (Figure 6) 65% of the PEs were identical to those of coarse-scale analysis, with 26% resulting from splitting and 9% of from lumping. Likewise, ES2 analyses yielded more PEs than fine-scale ES1 analyses, with 80 to 90% of PEs either identical to those of fine-scale ES1 analyses or resulting from splitting, with 2 to 17% resulting from lumping.

#### 3.2.1 Distribution Patterns Observed in Ecotype Simulation 1 Analyses

We first repeated ES1 analyses based on the HFS_50_ diversity in the neighbor-joining tree performed by Becraft et al. [2] to allow comparison of ES1 and ES2. Predominant PEs B′9, A1, A4 and A14 were predicted in coarse-scale analysis (orange lines in Figure 5). These PEs were previously found to be vertically stratified from top to bottom in the upper ∼1 mm-thick photic zone of the mat in the order listed (Supplemental Figure 6). The fine-scale analysis yielded nearly the same PEs, except some PEs were split into multiple PEs (red lines in Figure 5).

We used CCA analyses to test the predictions of the Stable Ecotype Model with the distribution of PEs in the CCA ordination space. Coarse-scale ES1 analyses of Becraft et al. [2] are presented in Supplemental Figure 6, but here we focus on the fine-scale ES1 analysis. The graphical presentation of PEs in the ordination space reflect their vertical stratification in the mat environment (Figure 7A), with top to bottom progression of PEs B′9, A1, A4, and A14. Note the distribution of centroids for each PE along the depth vector. These PEs yielded distributions that largely did not overlap, demonstrating ecological distinctness among PEs. We also used CCA to test whether each PE was ecologically homogeneous. The criterion for homogeneity was that the members of a PE should cluster together, non-randomly, in the environment represented by the CCA ordination space. All of these PEs (excluding PE A14) clustered, non-randomly, in the environment, suggesting that members within a given PE share similar ecological requirements and are thus interchangeable with each other. PEs with *p* > 0.05 might be due to insufficient sampling of variants within a PE or to lumping of >1 PE (see example below).

**Figure 7:**
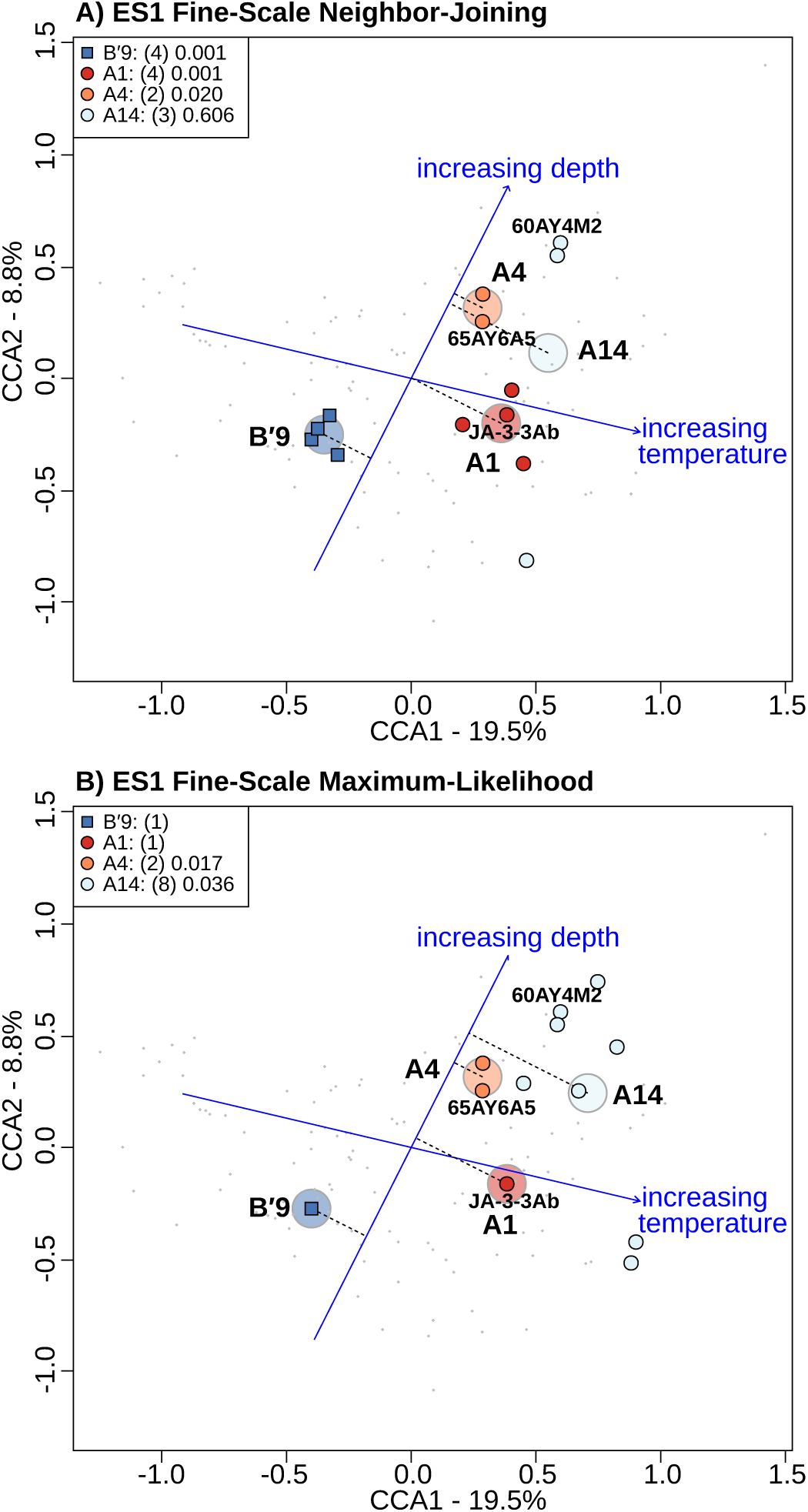
Canonical correspondence analysis highlighting *psaA* variants of predominant *Synechococcus* putative ecotypes (PEs) demarcated by the Ecotype Simulation 1 fine-scale method using (A) neighbor-joining and (B) maximum-likelihood phylogenies. Phylogenies were created from environmental *psaA* segments that numbered >50 (HFS_50_). Demarcation of PEs in A performed by Becraft et al. [2]. Large, lighter colored circles represent the centroids of highlighted PEs. Dotted lines connect each of these centroids to the depth vector to aid visualization of the distribution of PEs along the depth gradient measured. *Synechococcus* strains JA-3-3Ab, 65AY6A5, and 60AY4M2 share *psaA* sequences with HFS_50_ in these predominant PEs and are labeled on each plot. P-values are associated with the hypothesis that the members of PEs should not be distributed randomly.

Since ES2 uses a maximum-likelihood phylogeny, we next based our ES1 analyses on the HFS_50_ diversity in the maximum-likelihood tree, which presented a highly similar though not identical view of the A/B′-lineage *Synechococcus* phylogeny to that provided by a neighbor-joining phylogeny (compare Figures 5 and 6). The same predominant PEs detected in the fine-scale neighbor-joining analyses (PEs B′9, A1, A4 and A14) were detected in fine-scale maximum-likelihood analysis (orange and red lines in Figure 6B). In CCA analyses, these PEs exhibited the same vertical stratification from top to bottom in the upper ∼1 mm-thick photic zone of the mat, in the order listed above (Figure 7B), demonstrating ecological distinctness. The membership of each predominant PE that contained more than one HFS_50_ member (e.g., A4 and A14) clustered together, non-randomly, in the environment, demonstrating ecological interchangeability within each of these PEs. Similar results were found for coarse-scale analyses based on both treeing algorithms (compare Supplemental Figures 6A and 6B).

#### 3.2.2 Distribution Patterns Observed in Ecotype Simulation 2 Analyses

In the main text, we report results of ES2 analyses based on maximum-likelihood phylogenies. Results based on neighbor-joining phylogenies, which were highly similar, can be found in the Supplemental Information (Supplemental Figure 7). Analyses based on the HFS_50_ diversity in the maximum-likelihood tree detected predominant PEB31 (B′9), PEA20 (A1), PEA16 (A4), and PEA19 (A14) (blue lines Figure 6). In this analysis, there was a single PEB31 (B′9) variant and a single PEA20 (A1) variant, but PEA16 (A4) and PEA19 (A14) had multiple variants that clustered, non-randomly, in the environment analyzed with CCA (Figure 8A). Again, the top to bottom distribution was in the order observed above, PEB31 (B′9), PEA20 (A1), PEA16 (A4), and PEA19 (A14).

**Figure 8:**
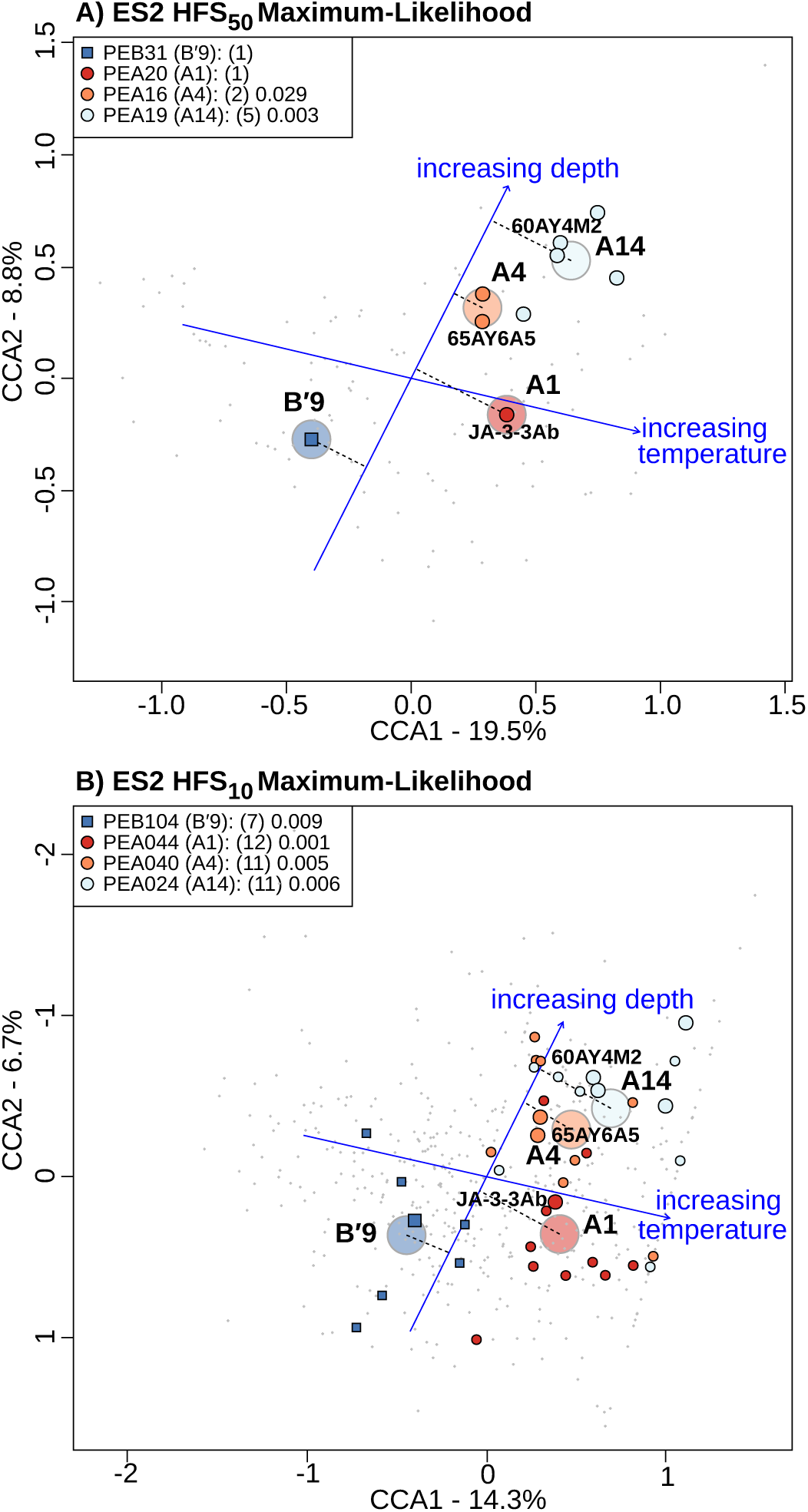
Canonical correspondence analysis highlighting *psaA* variants of predominant *Synechococcus* putative ecotypes (PEs) demarcated by Ecotype Simulation 2 using a maximum-likelihood phylogeny. Phylogenies were created from environmental *psaA* segments that numbered (A) >50 (HFS_50_), or (B) >10 (HFS_10_). Larger glyphs in B represent HFS_50_ environmental sequences, while smaller glyphs represent HFS_10_ environmental sequences. Large, lighter colored circles represent the centroids of highlighted PEs. Dotted lines connect each of these centroids to the depth vector to aid visualization of the distribution of PEs along the depth gradient measured. *Synechococcus* strains JA-3-3Ab, 65AY6A5, and 60AY4M2 share *psaA* sequences with HFS_50_ in these predominant PEs and are labeled on each plot. P-values are associated with the hypothesis that the members of PEs should not be distributed randomly.

ES2 analysis of HFS_10_ diversity, which could not be done with ES1 due to its excessive memory and CPU usage, increased the number of variants detected in each predominant PE simply due to the increased number of variants analyzed. ES2 analysis of HFS_10_ diversity in the maximum-likelihood tree resulted in 7-12 variants per PE. All predominant PEs exhibited non-random distributions in CCA analysis and the same vertical patterning previously observed (Figure 8B).

### 3.3 Comparison of ES1 and ES2 PE Predictions

We infer from these observations that ES2 demarcates PEs in a manner similar to that of ES1. While there are multiple differences between versions of the program that mainly affect the variants that are predicted to be within a PE, both analyses of the same dataset yielded similar PE clusters that distributed with depth in the mat similarly to that seen by Becraft et al. [2]. Furthermore, ES2 analysis enabled deeper sequence coverage, which improved the significance of non-randomness of most observed clusters. Demarcations based on maximum-likelihood phylogeny appeared to yield the best evidence of the existence and environmental distribution of different closely related *Synechococcus* PEs.

### 3.4 Analysis of the Full HFS_10_ Dataset

Analysis of the entire HFS_10_ dataset yielded a much larger number of PEs than those described here, though many are rare contributors to the dataset. These rare PEs are likely to represent rare members of the mat community (note the blue abundance bars in Figure 9). Significant environmental clustering in CCA (*p <* 0.05) is observed more frequently in the most abundant PEs (e.g., PEs A1, A4, A14; compare the orange p-value bars and the dashed red confidence interval lines with the corresponding blue bars in Figure 9). One exception is PE A6 of Becraft et al. [2], which is best explained by the observation that the partial *psaA* sequence used to demarcate species is shared by two widely divergent phylogenetic groups [33]. In most cases, however, partial *psaA* sequences permitted the detection of distinct ecological species. If horizontal gene exchange had been rampant, one would expect more well-sampled PE clusters to not be significantly clustered in CCA. The lower degree of significant clustering of variants in rare PEs may indicate the randomization of distributions, possibly due to these members being dispersed randomly into the community, as opposed to occupying a discrete niche within the community.

**Figure 9:**
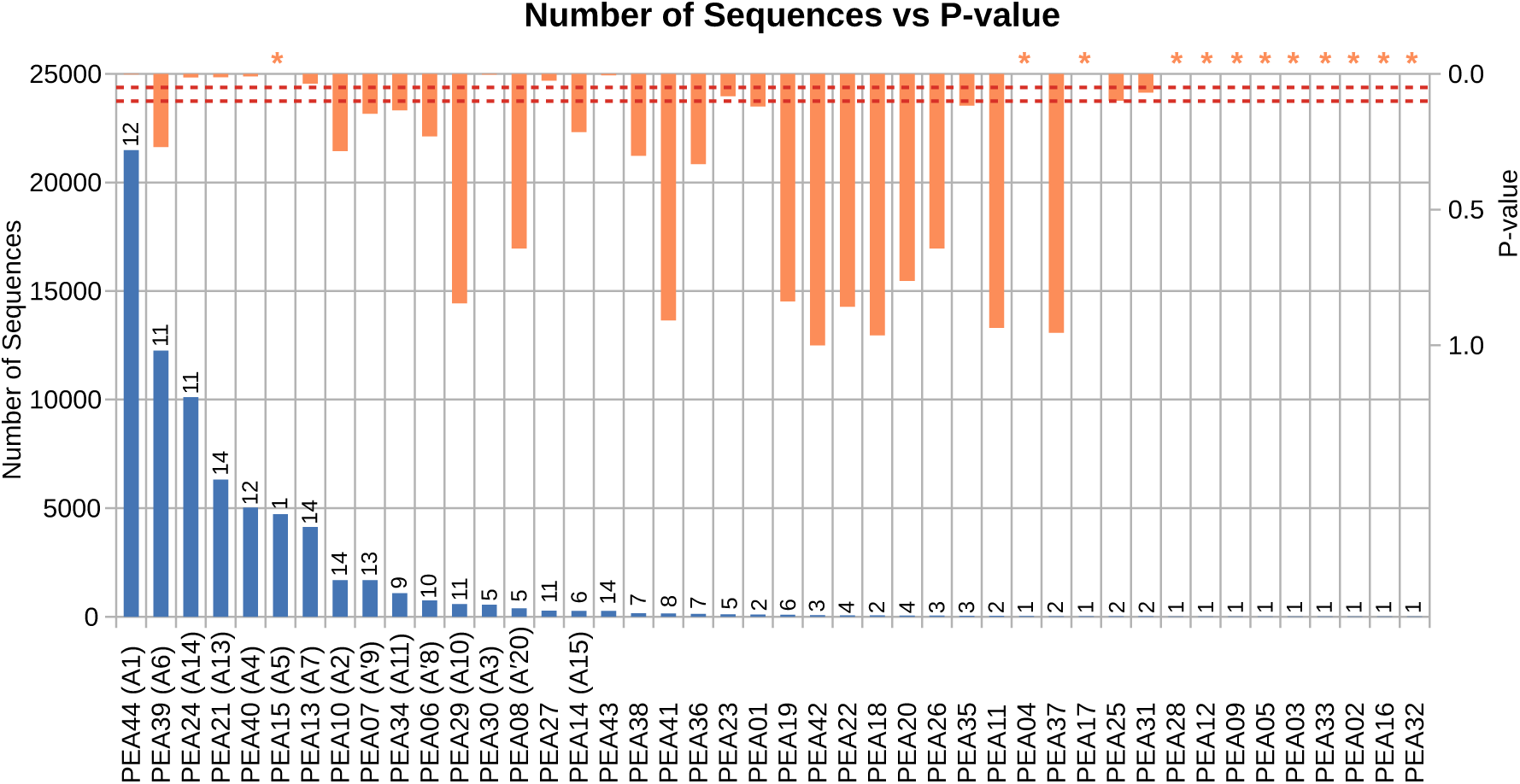
Sequence abundance of A-like *Synechococcus* HFS_10_ environmental *psaA* segments in putative ecotypes (PEs) demarcated by Ecotype Simulation 2 from the maximum-likelihood tree (blue bars). The number of unique sequence types found in each PE is included above the blue bars. Orange bars indicate p-values associated with the hypotheses that the members of PEs should not be distributed randomly. Orange asterisks above some columns represent p-values that could not be calculated since the PEs contained only a single member. The red dashed lines represent 0.05 and 0.10 confidence limits.

In an attempt to minimize the effect of a large variance in PE abundances, abundance counts were transformed using the formula: *n*_*transformed*_ = *log*(*n* + 1). Results of CCA analyses (not shown) where highly similar to the untransformed results presented in the manuscript. Significant environmental clustering in CCA (*p <* 0.05) is observed more frequently in the most abundant PEs.

### 3.5 Testing the Runtime of ES2

In order to test how well the changes made to ES2 were to meeting the goal of analyzing a large number of sequences, 191,901 unique A-like *Synechococcus* environmental sequences collected by Wood [44] were randomly subsampled to test the new algorithm on variously sized datasets. ES2 was used on 38 subsamples, with sample sizes ranging from 5,000 to 190,000 sequences, to predict PEs (Figure 10 and Supplemental Figure 5 for results from B′-like sequences). Although the maximum number of environmental sequences analyzed in this experiment (*n* = 190, 000) does not fully test the power of ES2, the overall linear nature of the runtime curve (see the blue line in Figure 10) and the fact that the average length of the analysis didn’t exceed six hours, illustrated that analyses of even larger datasets is possible. Such analyses would not be possible with ES1.

**Figure 10:**
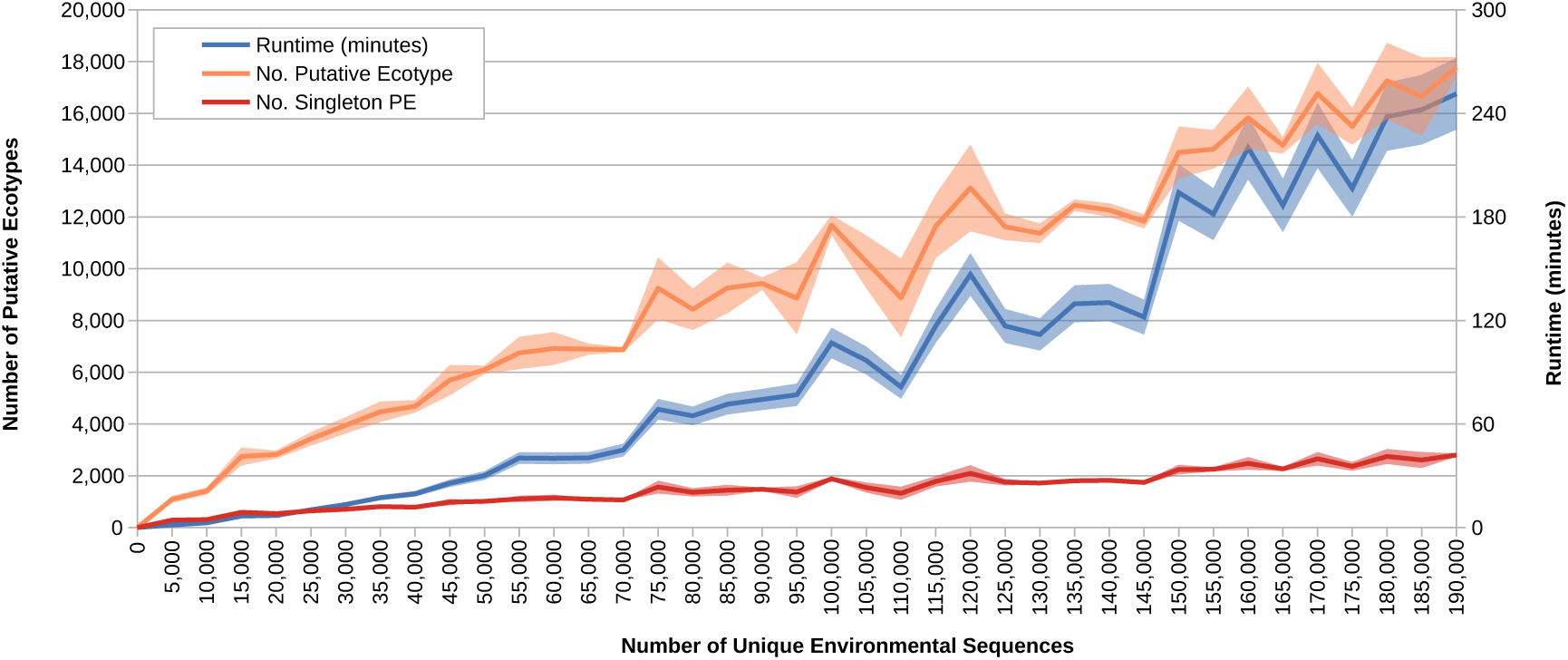
191,901 unique A-like *Synechococcus psaA* segments present in an environmental dataset containing 2,197,037 A-like *Synechococcus psaA* segments were subsampled at 5,000 sequence intervals. The darker colored lines represent the average and the shaded regions represent the standard error of three trials. The blue line and shaded region demonstrate the amount of time used to run the trial subsample using ES2 (on an Intel Core i7-6700 processor). The orange line and shaded region note the total number of PEs demarcated while the red line and shaded region note the number of PEs with only one member.

## 4 Conclusions

Ecotype demarcation results produced with ES2 are similar to those produced with ES1 with some cases of differential splitting or lumping that reflect the various changes in the treeing and simulation algorithms. Most importantly, these changes allow for the rapid analysis of large sequence datasets, which will permit researchers to explore fine-scale microbial diversity. Analyses based on HFS_10_ provide a clear example of the benefit of deeper coverage of sequence diversity. Since the same snapshot of the phylogeny is used at all stages of ES2, and FastTree is now incorporated for generating that phylogeny, we expect ES2 to provide a higher level of accuracy in ecotype demarcation than ES1 provided.

Although this study only examined the diversity within A/B′-lineage *Synechococcus* populations, we believe that ES2 will be applicable on a large scale because a) ES2 utilizes a theory-based model of speciation that functions on neutral genetic diversity present in the gene segment, gene, or suite of concatenated genes analyzed by the program, b) ES1 was shown to predict ecologically distinct ecotypes of *Bacillus* [21, 9] and ES2 performs similarly to ES1, and c) because it has been able to predict ecologically distinct PE with ecologically interchangeable members within other phototrophic taxa living in same system studied here [44].

Ecotype Simulation 2 is released under version 2 of the GNU General Public License and can be downloaded for Windows, Linux, and OSX from https://github.com/sandain/ecosim.

## Availability and requirements

**Project name:** Ecotype Simulation

**Project home page:** https://github.com/sandain/ecosim

**Operating systems:** Windows, OSX, Linux

**Programming languages:** Java, Fortran

**Other requirements:** Java 8 or higher

**License:** GNU GPL

## List of abbreviations

CCA: Canonical correspondence analysis
ES1: Ecotype Simulation version 1.
ES2: Ecotype Simulation version 2.
HFS: High Frequency Sequence (includes the minimum sequence count cutoff in subscript if applicable)
PE: Putative Ecotype.
npop: The estimated number of ecotype populations.
omega: Rate of net ecotype formation.
sigma: Rate of periodic selection.

## Declarations

### Ethics approval and consent to participate

Not applicable.

### Consent for publication

Not applicable.

### Availability of data and material

The sequence data analyzed for the HFS_50_ and HFS_10_ data sets used in this study are the same analyzed by Becraft et al. [2] and are available from MG-RAST (http://metagenomics.anl.gov; 4613896.3–4614007.3). The sequence data analyzed by Wood et al. [44] and utilized in this study to test ES2 with large datasets are available with the source code at https://github.com/sandain/ecosim.

### Author’s contributions

JW helped plan and develop ES2, analyzed barcode sequence data, performed canonical correspondence analyses and comparisons between algorithms, and assisted in preparation of the manuscript. EB tested early versions of ES2 and participated in preparation of the manuscript. DK helped plan and develop ES2 and participated in preparation of the manuscript. FC helped plan and develop ES2, co-supervised the research, and participated in preparation of the manuscript. DW obtained funding for the project, supervised the work, and participated in preparation of the manuscript.

## Acknowledgements

This study was conducted under Yellowstone National Park research permits YELL-0129 and 5494 (DW), and we appreciate the assistance from National Park Service personnel. We would also like to thank Lingyuan Ke, a student of DK, with help identifying the issue with using the Fortran 90 intrinsic pseudo-random number generator in a threaded environment.

