## Supplemental Information for "Ecotype Simulation 2: An improved algorithm for efficiently demarcating microbial species from large sequence datasets"

### 1 Analysis Of B′ *Synechococcus* Lineage

#### 1.1 Methods

B′-like *Synechococcus* high-frequency sequences (HFSs) were treated exactly like the A-like HFSs presented in the main text.

#### 1.2 Results And Discussion

As with the A-lineage presented in the main text, binning results for the B′-lineage are most similar between the Ecotype Simulation 1 (ES1) matrix-based method and the Ecotype Simulation 2 (ES2) result generated with a neighbor-joining tree (Supplemental Figure 1). In Supplemental Figures 2 and 3 PEs are numbered according to ES2 output, and correspondence with predominant PEs predicted using ES1 by Becraft et al. (2015) is shown in parentheses. As previously noted in that study, fine-sale ES1 analysis yielded a greater number of PEs than coarse-scale analysis, but this was mainly due to splitting coarse-scale PEs. for instance, in the fine-scale analyses shown in Supplemental Figures 2 and 3, all of the PEs in fine-scale analyses were either identical to those in coarse-scale analysis or resulted from splitting. ES2 analysis also yielded a greater number of PEs than found in coarse-scale ES1 anayses, and 88 to 95% of these were identical to ES1 coarse-scale PEs or splitting them into multiple PEs.

As with the A-lineage, analysis of the entire B′-lineage $HFS_{10}$ dataset yielded a much larger number of PEs than those described here (Supplemental Figure 4), though most were rare contributors to the dataset and are likely to be rare members of the mat community. Significant clustering in canonical correspondence analysis (CCA) ordination space was observed in highly abundant PEs PEB104 (B′9) and PEB105, and less abundant PEs PEB036 (B′7) and PEB023 (B′8-5) (Supplemental Figure 4). Other highly abundant B′-like PEs either didn't have enough HFS members to analyze with CCA (PE PEB079), or displayed disjoint distributions (i.e., $p > 0.05$). The B′-lineage has been hypothesized to be older, have more sequence diversity, and exhibits more evidence of recombination than the A-lineage (Melendrez et al., 2016), and as mentioned in the main text, recombination between distantly related PEs might explain the higher frequency of disjointly distributed B′-like PEs.

As with the runtime curve produced for the A-lineage in the main text, ES2 was able to predict ecotype membership for the B′-lineage in about six hours when analyzing 195,153 unique sequences sampled from a dataset containing 1,774,277 sequences (see the blue line in Supplemental Figure 5). The runtime curve was nearly linear with the number of environmental sequences analyzed during this test, illustrating that analyses of even larger datasets is possible.

### 2 ES1 Coarse-Scale Analyses Of Distribution Patterns

#### 2.1 Methods

Analyses of distribution patterns observed in ES1 coarse-scale analyses were performed using CCA as was done in the main text for ES1 fine-scale analyses.

#### 2.2 Results And Discussion

Similar to that found in fine-scale analyses presented in the main text, PEs are vertically stratified in the mat environment, with top to bottom progression of PEs B′9, A1, A4, and A14 using a neighbor-joining phylogeny (Supplemental Figure 6A), and PEs B′9, A4, and A1 using a maximum-likelihood phylogeny (Supplemental Figure 6B). Note that in coarse-scale maximum-likelihood analysis, PE A1 was lumped with PE A14 variants, explaining why there is no PE A14 in and possibly causing the deeper apparent distribution of PE A1. As was noted in fine-scale analyses, the coarse-scale PEs yielded distribution patterns that largely did not overlap, again demonstrating the ecological distinctness of these PEs. All of these PEs (excluding A14 in neighbor-joining analysis) clustered, non-randomly, in the environment, suggesting PE members share similar ecological requirements.

### 3 ES2 Analyses Based On Neighbor-Joining Phylogenies

#### 3.1 Methods

ES2 analyses were based on neighbor-joining phylogenies described in the main text. Analyses were performed on phylogenies generated from both the $HFS_{50}$ and $HFS_{10}$ datasets.

#### 3.2 Results And Discussion

ES2 analyses based on the $HFS_{50}$ diversity in the neighbor-joining tree detected ecotype diversity in the B′9 clade, with PEB05 (B′9) sharing the same $HFS_{50}$ that were detected in ES1 fine-scale analysis, but PEB01 contained fewer variants (blue lines in main text Figure 5). The PEB05 (B′9) variants formed a discrete, non-randomly distributed cluster in CCA analyses (Supplemental Figure 7A), and was distributed higher in the mat. ES2 analysis split PEs A1, A4 and A14 into 2-3 PEs each. In each of these PEs, only a single variant matched the variants previously used to define them, so that clustering could not be analyzed. However, the distribution of these PEs was consistent with that observed in ES1 analyses (Supplemental Figure 7A), with the PEA13 (A1) variant above the PEA07 (A4) variant, which was above the PEA19 (A14) variant.

Analysis based on the $HFS_{10}$ diversity in the neighbor-joining tree resulted in 2-9 variants per PE. $HFS_{10}$ variants in PEB076 (B′9), and PEA083 (A1) clustered non-randomly in the environment analyzed with CCA (Supplemental Figure 7B). PEA030 (A4) exhibited 4 variants, which appeared to possibly form a non-random cluster ($P = 0.091$). PEA069 (A14) exhibited 2 variants whose clustering could not be shown to be significantly nonrandom ($P = 0.271$).

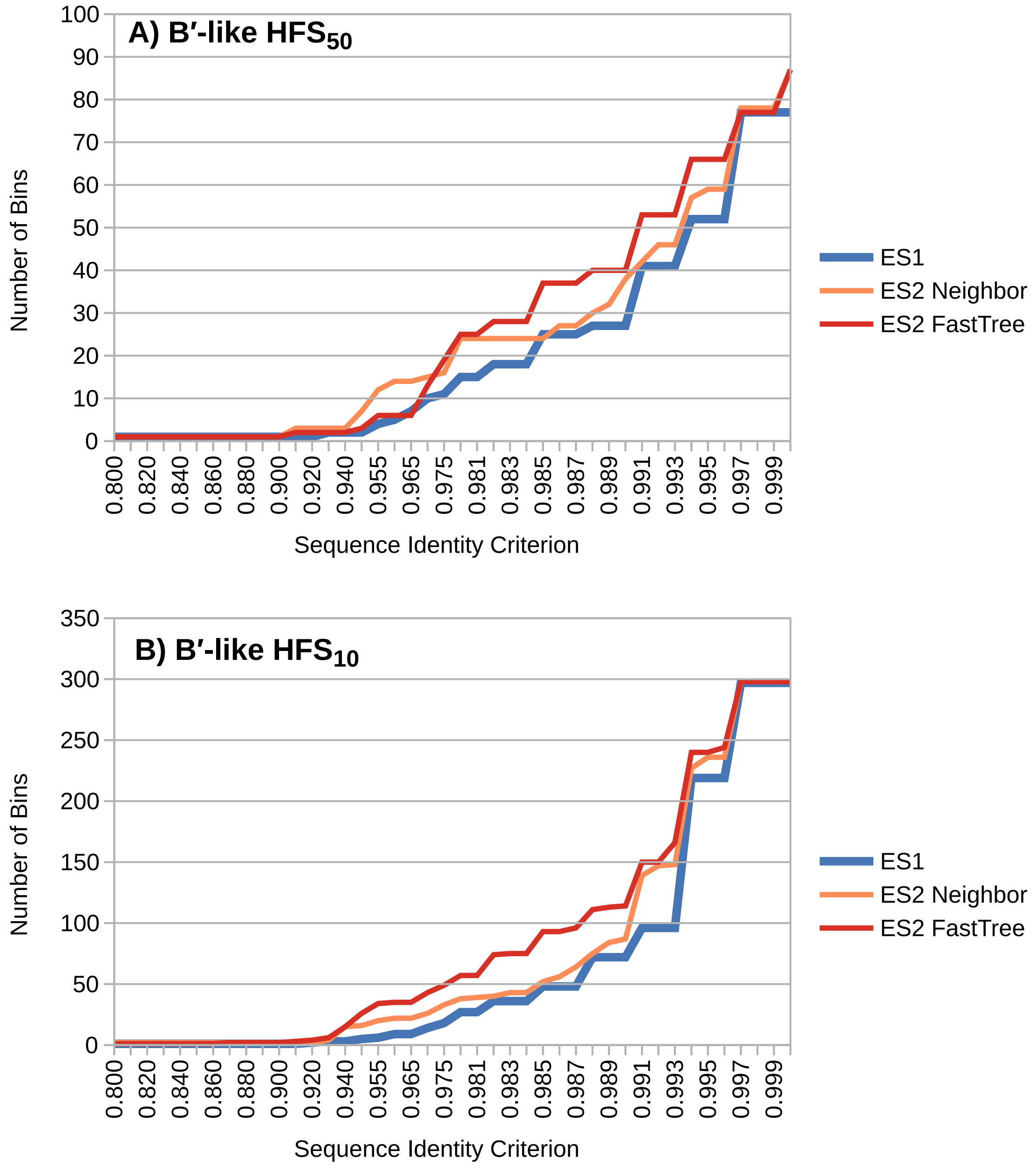

Supplemental Figure 1: Binning results using B′-like *Synechococcus psaA* high-frequency sequences (HFS) occuring at least (A) fifty times ($HFS_{50}$) and (B) ten times ($HFS_{10}$) in the environmental sequence dataset.

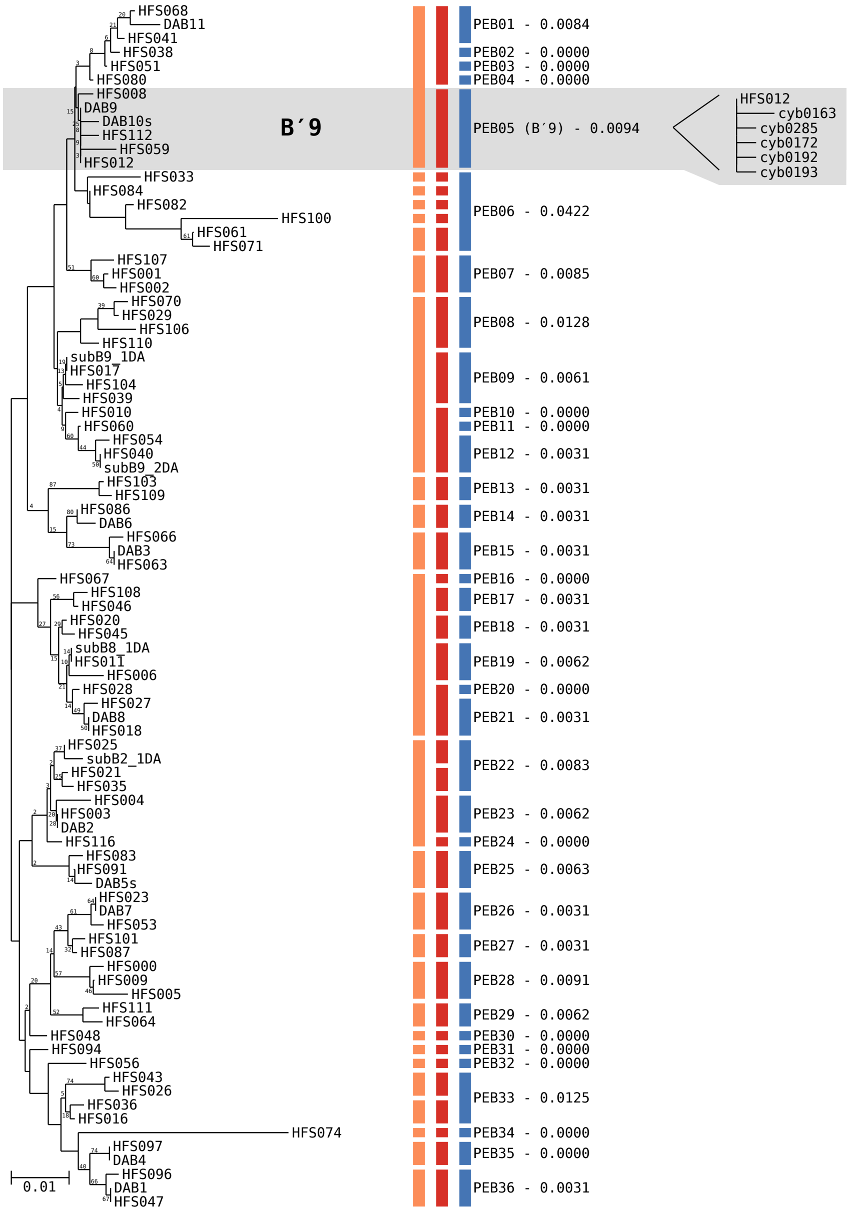

Supplemental Figure 2: Neighbor-joining phylogeny with putative ecotype (PE) demarcation of B′-like *Synechococcus* HFS$_{50}$ environmental *psaA* segments generated using PHYLIP. Gray shading denotes predominant PE B′9 demarcated using Ecotype Simulation 1 (ES1) that is examined in detail in the main text. Colored vertical bars indicate demarcation done by different algorithms, from left to right: orange, ES1 coarse-scale; red, ES1 fine-scale; and blue, Ecotype Simulation 2 (ES2). PEs are labeled with the ES2 demarcation and include the maximum distance among members of the clade. ES2-demarcated PEs that contain the same dominant variant as a PE demarcated by Becraft et al. (2015) using ES1 are indicated in parentheses after the ES2-generated PE names. The portion of the HFS$_{10}$ neighbor-joining tree corresponding to predominant B′-like PE PEB05 (B′9) is included on the right for comparison. The scale bar represent 0.01 nucleotide substitutions per site.

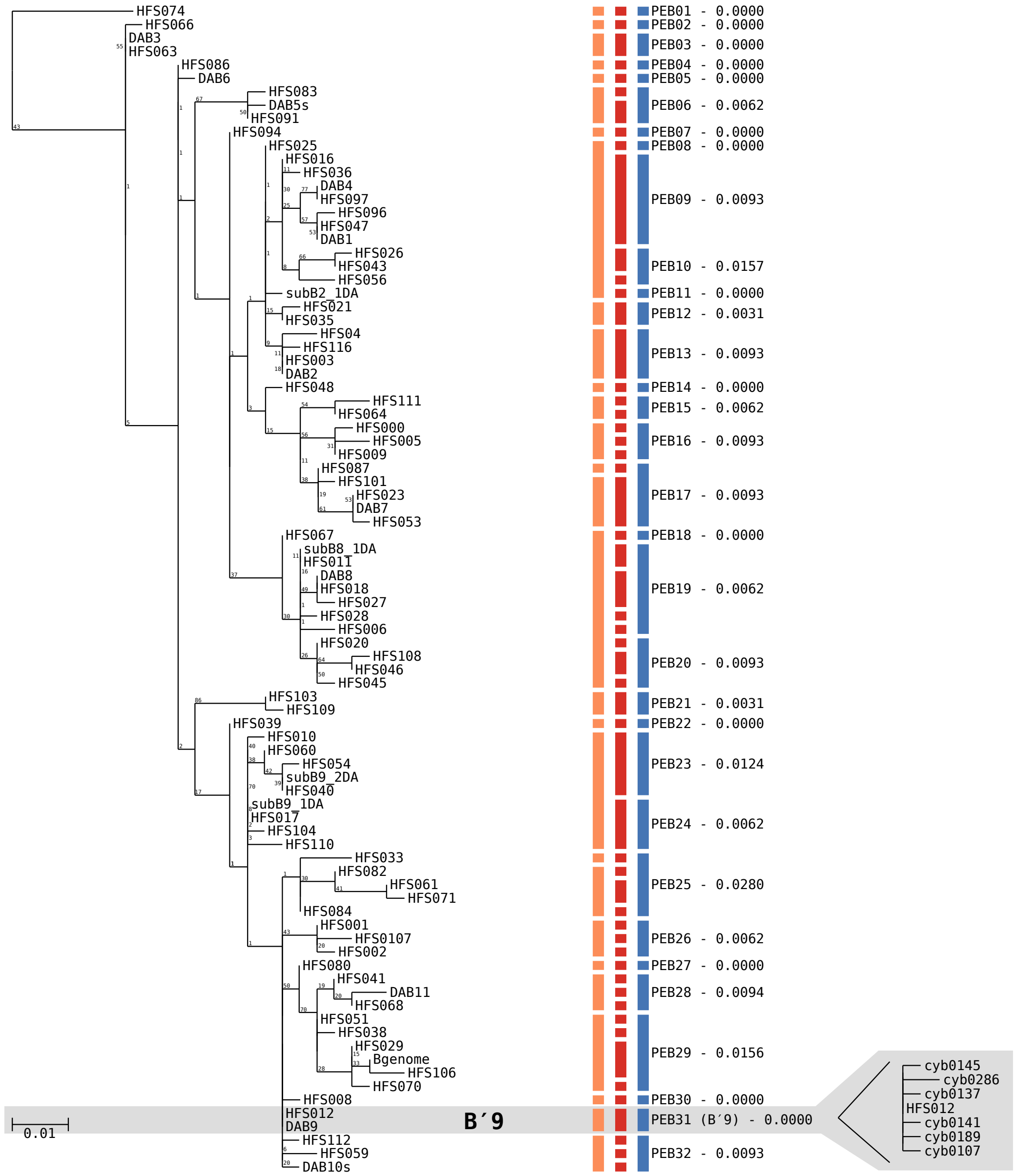

Supplemental Figure 3: Maximum-likelihood phylogeny with putative ecotype (PE) demarcation of B′-like *Synechococcus* $HFS_{50}$ environmental *psaA* segments generated using FastTree. Gray shading denotes predominant PE B′9 demarcated using Ecotype Simulation 1 (ES1) that is examined in detail in the main text. Colored bars indicate demarcation done by different algorithms, from left to right: orange, ES1 coarse-scale; red, ES1 fine-scale; and blue, Ecotype Simulation 2 (ES2). PEs are labeled with the ES2 demarcation and include the maximum distance among members of the clade. ES2-demarcated PEs that contain the same dominant variant as a PE demarcated by Becraft et al. (2015) using ES1 are indicated in parentheses after the ES2-generated PE names. The portion of the $HFS_{10}$ maximum-likelihood tree corresponding to predominant B′-like PE PEB31 (B′9) is included on the right for comparison. The scale bar represent 0.01 nucleotide substitutions per site.

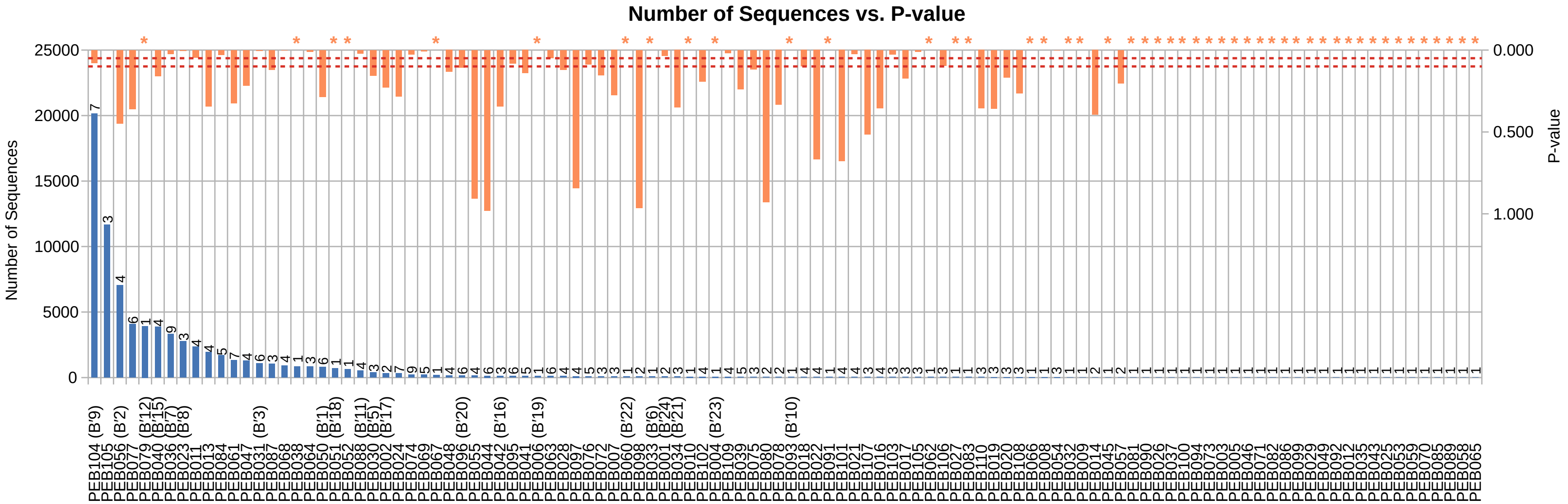

Supplemental Figure 4: Sequence abundance of B′-like *Synechococcus* $HFS_{10}$ environmental *psaA* segments in putative ecotypes (PEs) demarcated by Ecotype Simulation 2 from the maximum-likelihood tree (blue bars). The number of sequence types found in each PE is included above the blue bars. Orange bars indicate p-values associated with the hypotheses that the members of PEs should not be distributed randomly. Orange asterisks above some columns represent p-values that could not be calculated since the PEs contained only a single member. The red dashed lines represent 0.05 and 0.10 confidence limits.

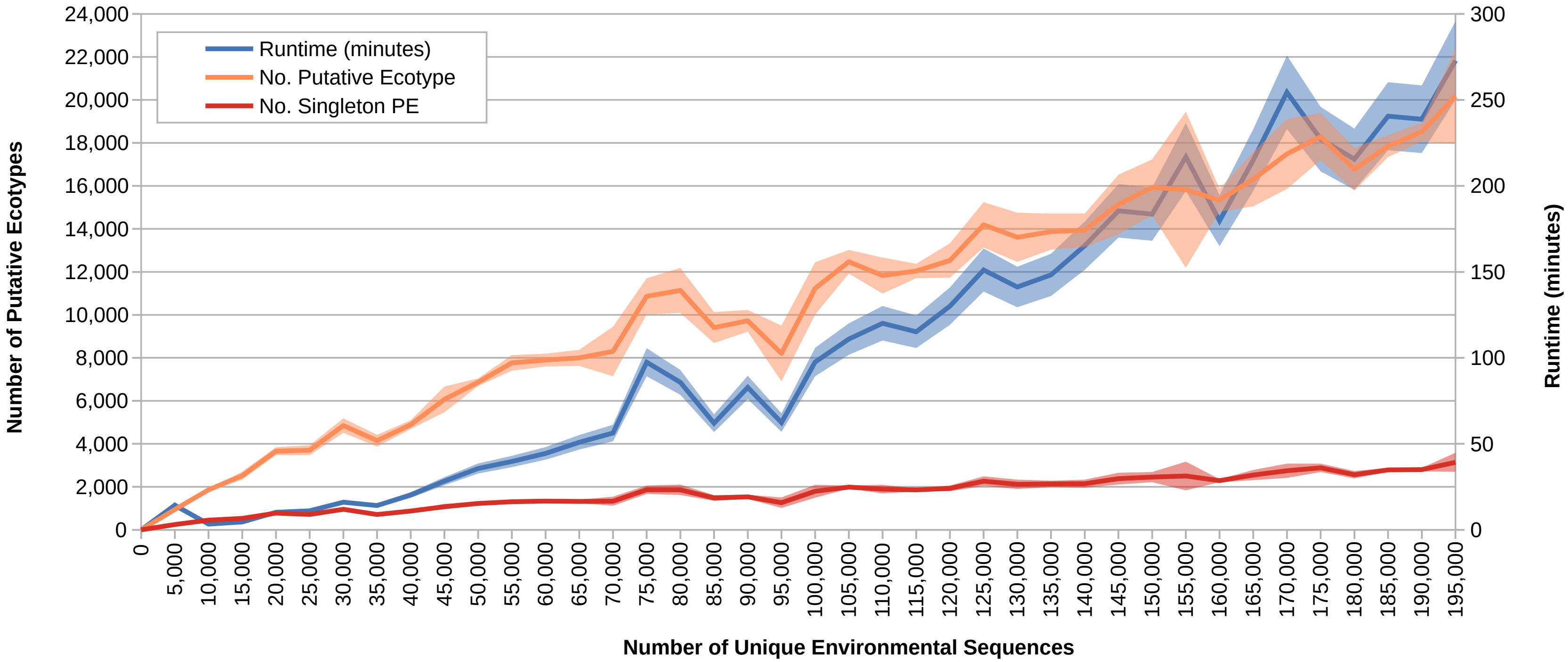

Supplemental Figure 5: Number of putative ecotypes (PEs) demarcated and runtime of repeated subsampling of 195,153 unique B′-like *Synechococcus psaA* segments present in an environmental dataset containing 1,774,277 B′-like *Synechococcus psaA* segments, with subsequent analysis by Ecotype Simulation 2 (ES2). The darker colored lines represent the average and the shaded regions represent the standard error of three trials. The blue line and shaded region demonstrate the amount of time used to run the trial subsample using ES2 (on an Intel Core i7-6700 processor). The orange line and shaded region note the total number of PEs demarcated while the red line and shaded region note the number of PEs with only one member.

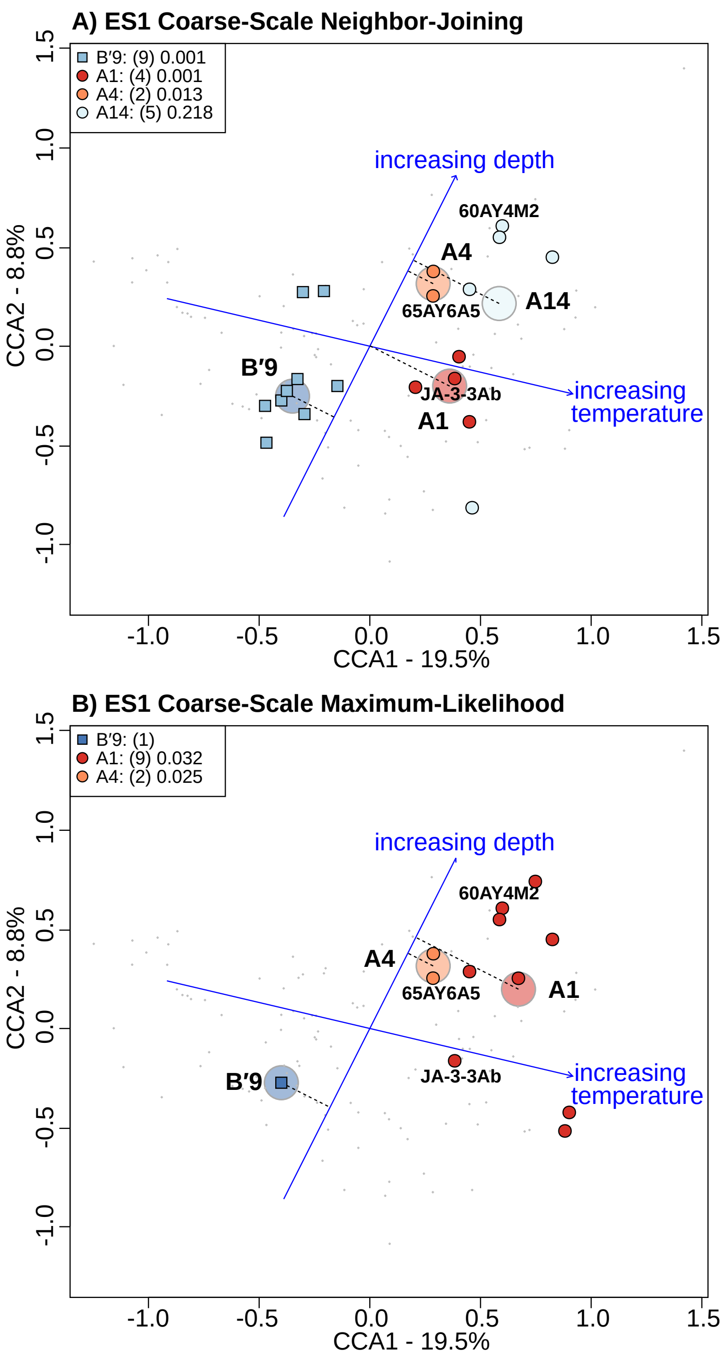

Supplemental Figure 6: Canonical correspondence analysis highlighting *psaA* variants of predominant *Synechococcus* putative ecotypes (PEs) demarcated by the Ecotype Simulation 1 coarse-scale method using (A) neighbor-joining and (B) maximum-likelihood phylogenies. Phylogenies were created from environmental *psaA* segments that numbered >50 ($HFS_{50}$). Demarcation of PEs in A performed by Becraft et al. (2015). Large, lighter colored circles represent the centroids of highlighted PEs. Dotted lines connect each of these centroids to the depth vector to aid visualization of the distribution of PEs along the depth gradient measured. *Synechococcus* strains JA-3-3Ab, 65AY6A5, and 60AY4M2 share *psaA* sequences with $HFS_{50}$ in these predominant PEs and are labeled on each plot. P-values are associated with the hypothesis that the members of PEs should not be distributed randomly.

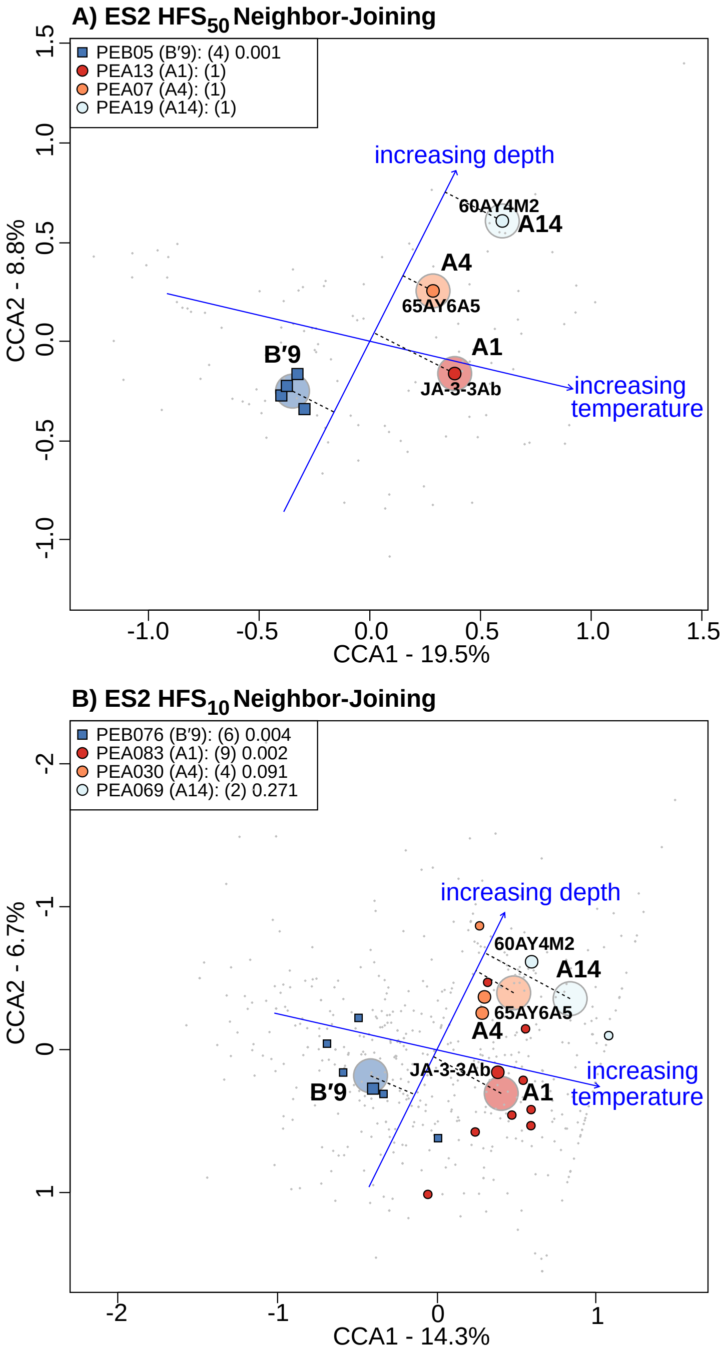

Supplemental Figure 7: Canonical correspondence analysis highlighting *psaA* variants of predominant *Synechococcus* putative ecotypes (PEs) demarcated by Ecotype Simulation 2 using a neighbor-joining phylogeny. Phylogenies were created from environmental *psaA* segments that numbered (A) >50 ($HFS_{50}$), or (B) >10 ($HFS_{10}$). Larger glyphs in B represent $HFS_{50}$ environmental sequences, while smaller glyphs represent $HFS_{10}$ environmental sequences. Large, lighter colored circles represent the centroids of highlighted PEs. Dotted lines connect each of these centroids to the depth vector to aid visualization of the distribution of PEs along the depth gradient measured. *Synechococcus* strains JA-3-3Ab, 65AY6A5, and 60AY4M2 share *psaA* sequences with $HFS_{50}$ in these predominant PEs and are labeled on each plot. P-values are associated with the hypothesis that the members of PEs should not be distributed randomly.
